# Epistasis between synonymous and nonsynonymous mutations in *Dictyostelium discoideum* ammonium transporter *amtA* drives functional complementation in *Saccharomyces cerevisiae*

**DOI:** 10.1101/2022.04.05.486919

**Authors:** Asha Densi, Revathi S Iyer, Paike Jayadeva Bhat

## Abstract

Role of <u>H</u>orizontal <u>G</u>ene <u>T</u>ransfer (HGT) in evolution transcends across the three domains of life. Ammonium transporters are present in all species and therefore offer an excellent paradigm to study protein evolution following HGT. While investigating HGT through complementation assay, we observed that synonymous and nonsynonymous mutations follow an epistastic relationship. As a proxy for HGT, we attempted to complement a *mep1mep2mep3Δ* strain of *S. cerevisiae* (triple deletion strain) which cannot grow on ammonium as a sole nitrogen source below a concentration of 3 mM, with *amtA* of *D. discoideum*. As the wild type *amtA* did not complement, we isolated two mutant derivatives of *amtA* that complemented the triple deletion strain of *S. cerevisiae*. *amtA M1* bears three nonsynonymous and two synonymous substitutions and these substitutions are necessary for its functionality. *amtA M2* bears two nonsynonymous and one synonymous substitution, all of which are necessary for functionality. These mutants were then studied at phenotypic, cell biological, and biochemical level. Interestingly, AmtA M1 transports ammonium but does not confer toxicity to methylamine while AmtA M2 transports ammonium as well as confers methylamine toxicity, demonstrating functional diversification. Based on the results presented, we suggest that protein evolution cannot be fathomed by studying nonsynonymous and synonymous substitutions separately. This is because, protein evolution entails an interaction between synonymous and nonsynonymous substitution, which seems to have gone unnoticed thus far. Above observations have significant implications in various facets of biological processes and are discussed in detail.

**Highlights:**

- Ammonium transporters (Amts) from bacteria to humans complement in yeast
- AmtA of *D. discoideum* does not complement yeast defective for ammonium uptake
- Synonymous & nonsynonymous mutations are essential for AmtA functionality in yeast
- Conformational differences underlie functionality & functional diversification
- Protein evolution entails interaction of synonymous & nonsynonymous mutations

## Introduction

Current understanding of organic evolution is mainly shaped by the Darwinian concept of descent with modification, wherein, genetic material is vertically transmitted from parent to offspring [1]. In contrast, transfer of genetic material between two separate genomes, commonly referred to as horizontal gene transfer or lateral gene transfer, was initially observed in bacteria [2][3][4]. Therefore, the role of HGT in evolution is mainly studied in prokaryotes [5][6][7], with limited acceptance of its role in eucaryotes [8]. It is now abundantly clear that HGT can occur between prokaryotes and eucaryotes [8][9][10][11][12][13][14] and between eucaryotes themselves [15][16][17][18][19][20]. The phenomena of gene duplication and neofunctionalization [21], one of the main evolutionary forces considered early on under the paradigm of vertical gene transmission, have been shown to occur following HGT [22][23][24][25]. These developments bring in HGT well within the realms of Darwinian evolutionary thinking [26][27][28].

Despite the potential of rapid evolutionary changes [29], HGT is fraught with many barriers that are not mutually exclusive. For example, wide variation in the GC content [30][31][32], phylogenetic distance [33], codon adaptive index [31][34][35][36], increased fitness cost [37], and incompatibility in the biochemical mechanisms [38] are known to impede HGT [39]. The ‘complexity hypothesis’ posits that genes coding for standalone function such as, nutrient transport or metabolic functions are more prone for HGT [40][41] as compared to genes that code for informational molecules. Fungi, once thought to be recalcitrant for HGT because of their cell wall structure and osmotrophic feeding habits [42], are now known to have evolved even niche diversification because of HGT. For example, dihydroorotate dehydrogenase, an enzyme involved in pyrimidine biosynthesis, was acquired through HGT from *Lactobacillus* which conferred anaerobic life style in *Saccharomyces cerevisiae* [43][44][45]. Since then, many studies have reported HGT of genes involved in vitamin [46][47] and carbon utilization [48][49]. Genome sequence analysis of wine yeast revealed three distinct chromosomal regions consisting of genes that seems to have been acquired through HGT that provided an adaptive value for growth on conditions with high sugar and low nitrogen [50]. Despite these rapid technological advances in detecting HGT, our knowledge of the molecular events that allow foreign genetic material to get assimilated, fixed and functionally diversified in the recipient, is limited.

Ammonium transporters belong to Amt/Mep/Rh family, the members of which are distributed in all the three domains of life. Members of this family have undergone extensive duplication, sub- and neo-functionalisation, gene fusion and HGT across kingdoms [51]. For example, it has been proposed that few species of archaea, acquired Rh genes through HGT from eucaryotes [52]. An independent bioinformatic analysis indicated that members of Amt are vertically transmitted while members of Mep family have undergone extensive HGT [53][54]. In general, ammonium transporters from any one of the subfamilies are represented in most taxa except amoeba, nematods, which harbour members belonging to both Amt and Rh subfamily [55]. Members of Mep are present only in fungi, while vascular plants have only members of Amt subfamily [54]. Thus, the origin and the functional diversification of Meps in general is not clearly understood.

We carried out a complementation assay to simulate HGT between *D. discoideum* and *S. cerevisiae*, which are unicellular eucaryotes separated by 1480 million years of evolutionary divergence [56]. *D. discoideum* diverged from metazoan lineage after the plants but before yeast [57]. Ammonium transporter *amtA* of *D. discoideum*, an orthologue of *S. cerevisiae MEP2* [58], failed to complement *S. cerevisiae* strain lacking *MEP1, 2,* and *3*, for growth on ammonium (hereafter referring to the sum of NH_4_^+^ and NH_3_) below a concentration of 3 mM as the sole source of nitrogen. We isolated two independent mutants of *amtA* from two randomly generated mutant plasmid libraries. Surprisingly, we observed that in both mutants, a combination of synonymous as well as nonsynonymous mutations is required to confer the ammonium uptake activity to AmtA in *S. cerevisiae*. Detailed analysis indicated that synonymous and nonsynonymous mutations exhibit a clear epistatic relationship. These observations demonstrate the underlying molecular events that could possibly occur during HGT. These observations also challenge a long held notion that the ‘second site functional compensation’ (occurs when a loss of function mutation is suppressed by a second site mutation) in protein occurs only through nonsynonymous substitution.

## Results

### D. discoideum amtA does not complement S. cerevisiae triple deletion strain for growth on minimal medium containing ammonium below a concentration of 3mM

Functional complementation is normally used as a proxy to investigate the genetic relatedness between species and decipher structure function relationship of proteins. For example, ammonium transporters from humans [59], plants [60], bacteria [61], and worms [62] functionally complement *S. cerevisiae* triple deletion strain, which otherwise does not grow on synthetic medium containing ammonium at a concentration below 3mM. However, these complementation experiments do not throw much light on mechanisms of evolutionary changes during extensive HGT. To explore this, we decided to study the ability of ammonium transporter *amtA* of *D. discoideum* to complement a triple deletion strain of *S. cerevisiae*. *D. discoideum* is prone to HGT due to the fact that it is phagocytic and thus a good model to study. Further, the GC content, one of the known barriers of HGT, of these two organisms (22.4% in *D. discoideum* vs 38% in *S. cerevisiae*) is widely different. Therefore, it was imperative to look at the codon adaptive index (CAI) as well as the relative usage frequencies of the synonymous codons (%MinMax) of the ammonium transporters of these two organisms and compare it to the ammonium transporters of the organisms whose ammonium transporters are known to complement the *S. cerevisiae* triple deletion strain [63][64]. Surprisingly, CAI of *amtA* (*D. discoideum)* was higher than CAI of *RHAG* (*H. sapiens*), *AMT1-1* (*A. thaliana*), and *amtB* (*E. coli*) when compared to *S. cerevisiae* genome (Table 1). %MinMax values of *amtA* (*D. discoideum* vs. *S. cerevisiae)* was higher than *MEP2* (*S. cerevisiae* vs. *S. cerevisiae)*, while they were less for *RHAG* (*H. sapiens* vs. *S. cerevisiae*), *AMT1-1* (*A. thaliana* vs. *S. cerevisiae*), and *amtB* (*E. coli* vs. *S. cerevisiae*) (Figure 1A). This data suggested that *D. discoideum amtA* codons are more favourable for translation in *S. cerevisiae* as compared to *RHAG*, *AMT1-1*, *amtB*, and surprisingly to its own ammonium transporter *MEP2*. Based on these bioinformatics analyses we hypothesized that the difference in the GC content may not impede AmtA functionality in *S. cerevisiae*. This prompted us to explore the biological significance of codon usage in HGT using a simple complementation analysis.

**Figure 1:**
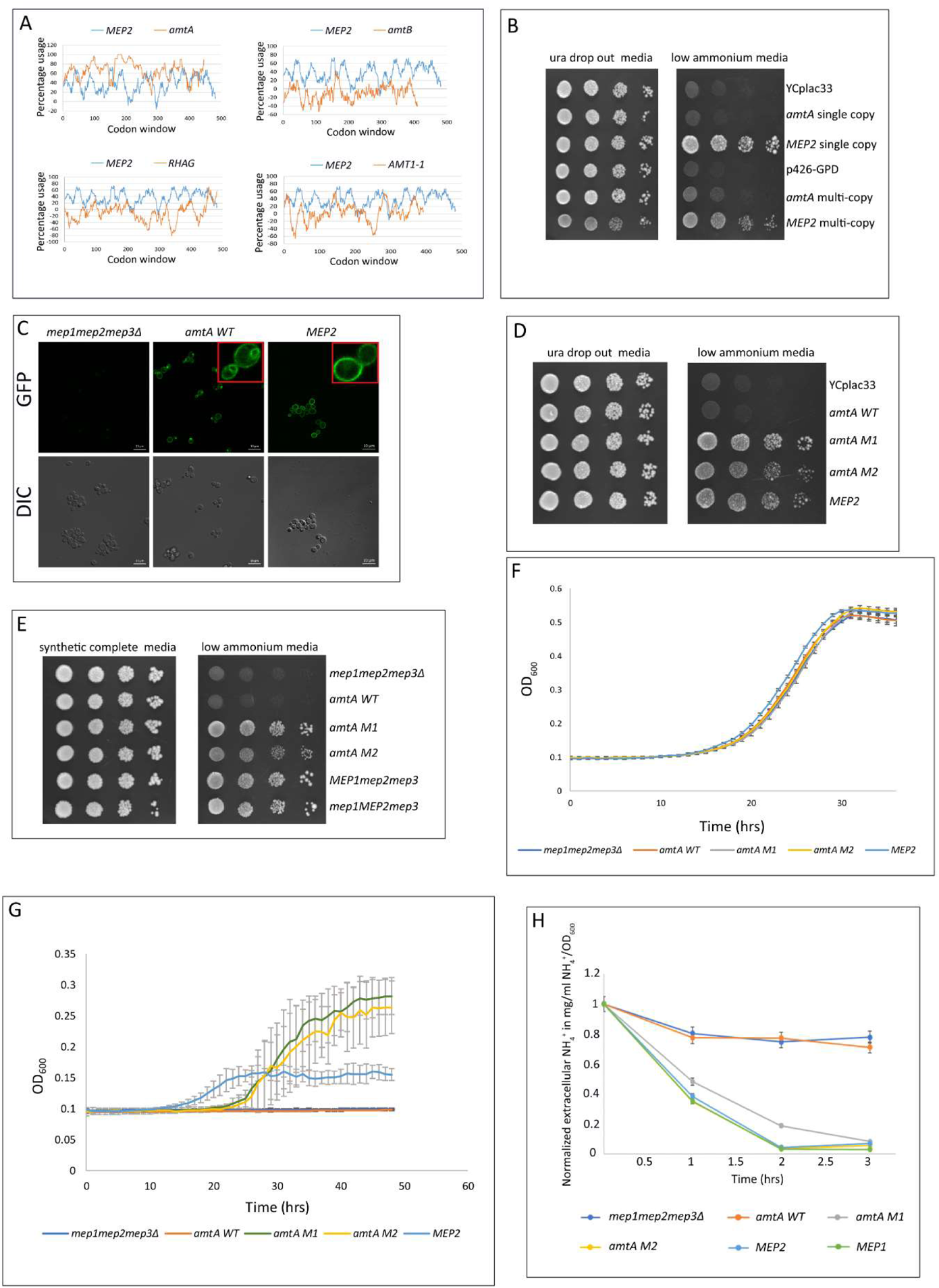
*amtA* mutants *M1* and *M2* but not *amtA WT* complement triple deletion strain. **(A) %MinMax of ammonium transporters expressed in *S. cerevisiae*.** %MinMax values of ammonium transporters when expressed in *S. cerevisiae* was obtained using %MinMax calculator online software [63][64]. Codon usage of *S. cerevisiae* was used. Percentage usage of rare codons is plotted again a sliding codon window of 18 codons. **(B) Growth of transformants on low ammonium**. 5 µl of the transformants of the triple deletion strain pre-grown in ura drop out glucose medium containing 20 mM ammonium to a cell density of 10^7^/ml were serially diluted and were spotted onto ura drop out glucose medium with 20 mM ammonium sulfate (left panel) and low ammonium media (right panel). The growth pattern was photographed after 4 days of incubation at 30°C. **(C) Expression and localization of yEGFP-tagged proteins.** Strains with yEGFP tagged proteins were grown in 0.1% proline media for 22 hours. The localization of the yEGFP tagged proteins was visualized using live fluorescence microscopy. The top panel represents the GFP fluorescence images, and the bottom panel represents DIC images. **(D) Growth of transformants on low ammonium.** 5 µl of the transformants of the triple deletion strain pre-grown in ura drop out glucose medium containing 20 mM ammonium to a cell density of 10^7^/ml were serially diluted and were spotted onto ura drop out glucose medium with 20 mM ammonium sulfate (left panel) and low ammonium media (right panel). The growth pattern was photographed after 4 days of incubation at 30°C. **(E) Growth of strains on low ammonium**. 5 µl of the strains pre-grown in synthetic complete glucose medium containing 20 mM ammonium to a cell density of 10^7^/ml were serially diluted and were spotted onto synthetic complete glucose medium with 20 mM ammonium sulfate (left panel) and low ammonium media (right panel). The growth pattern was photographed after 4 days of incubation at 30°C. **(F) Growth kinetics of strains on 0.1% proline media.** Different strains, pre-grown in 0.1% proline media, were grown in 0.1% proline media, and OD_600_ was measured as a function of time for 36 hours. **(G) Growth kinetics of strains on low ammonium.** Different strains, pre-grown in 0.1% proline media, were grown in low ammonium media, and OD_600_ was measured as a function of time for 48 hours. **(H) External ammonium concentration (in mg/ml NH_4_^+^/OD_600_) as a function of time.** Different strains were grown in 0.1% proline media till OD_600_ reaches 1.0. Cells were collected and transferred to a medium containing 0.1% proline and 500 µM ammonium at OD_600_ = 1.0. At each time point, cells were collected, filtered, and the filtrate estimated for ammonium as a function of time.

**Table 1:** CAI of different ammonium transporters when expressed in *S. cerevisiae*.

| **Ammonium transporter** | **CAI** |
| --- | --- |
| *D. discoideum AmtA* | 0.86 |
| *E. coli AmtB* | 0.58 |
| *A. thaliana Amt1-1* | 0.62 |
| *H. sapiens RhAG* | 0.67 |

Triple deletion strain was transformed with single (YCplac33) and multicopy (p426-GPD) plasmids bearing either *amtA* or *MEP2*, whose expression was driven by *MEP2* promoter. These transformants were pre-grown in uracil drop out media containing ammonium sulphate at a concentration of 20 mM, to a cell density of 5×10^7^/ml and serially diluted and spotted on uracil drop out media and minimal medium containing ammonium at a concentration of 500 µM as the sole nitrogen source (henceforth referred to as low ammonium media) (Figure 1B). Uracil dropout media is used to maintain the plasmids in the transformants. Transformant bearing the vector YCplac33 showed no growth while the positive control bearing either single or multicopy *MEP2* showed growth as expected. It was observed that triple deletion strain transformed with the multicopy *MEP2* consistently showed a slight growth retardation as compared to the transformant bearing single copy *MEP2* even on complete medium (Figure 1B). The transformants of triple deletion strain bearing single copy *amtA* or multicopy *amtA* did not grow on low ammonium media. The growth of these transformants were tested by streaking for single colonies on low ammonium media. No colonies were observed when the triple deletion strain transformed with either single or multicopy *amtA* were streaked for single colonies (Figure S1), clearly indicating that AmtA does not function as ammonium transporter in *S. cerevisiae*.

Above results were unexpected as orthologues of ammonium transporters from other evolutionarily distant species complement the triple deletion strain for growth on low ammonium [59][60][61][62]. Further, despite the codon usage of *D. discoideum* being favourable, *amtA* failed to complement the triple deletion strain. We wanted to ascertain that the incompatibility observed in our experiments is not due to a defect in expression and/or plasma membrane localization. To test this possibility *yEGFP* was fused to the C-terminus of *amtA* and was integrated at the *MEP2* locus such that the expression of *amtA::GFP* was driven by *MEP2* promoter. *yEGFP* was similarly integrated to the C-terminus of *MEP2*. These strains were grown in 0.1% proline, a poor nitrogen source for 22 hours. Under these growth conditions, *MEP2* promoter is transcriptionally active [65]. Live cells were observed under confocal microscope (Figure 1C). GFP signal could be clearly detected in the plasma membrane of *amtA::GFP* strain. In addition, GFP signal could also be detected in subcellular compartment in the cell. As expected, it can be observed that GFP signal was localized in the plasma membrane in strain expressing *MEP2::GFP* construct. Based on this, we inferred that the inability of *amtA* to complement a triple deletion yeast strain on low ammonium media may not be because of a defect in the expression or localization.

Based on the above, we hypothesized that most likely, the conformation of AmtA when expressed in *S. cerevisiae* is incompatible with the its normal function. It could be because of reasons such as aberrant co-translational folding leading to non-functional protein [66], or could be due to the difference in lipid composition of the membrane which can influence the conformation of proteins [67]. Substituting with frequently used codons to increase the translation rate, reducing GC content to reduce the possibility of stable mRNA secondary structures or even generating harmonised gene variant are some of the commonly used strategies to improve the expression/functionality of the heterologously expressed genes [68][69][70][71]. We reasoned that using the above approaches to obtain a functional protein, would not give us the insights in to the trajectory of protein evolution. Therefore, we resorted to a reverse genetic approach. We constructed two independent libraries in a single copy plasmid using two independent error prone PCR reactions, followed by ligation through in vivo recombination (Figure S2). Of the two, one set of reactions was carried out in presence of 0.35 mM MnCl_2_ while the other with 0.45 mM MnCl_2_. The expression of mutant ORFs was driven by *MEP2* promoter. We screened the above two libraries (0.35 mM MnCl_2_ and 0.45 mM MnCl_2_), each consisting of 25,000 colonies to identify transformants capable of growing on low ammonium media by replica plating. We obtained two transformants one from each library that grew on low ammonium media and these mutant clones are referred to as *amtA M1* and *amtA M2*. The plasmids from these transformants were isolated, and amplified in *E. coli*, and triple deletion strain was retransformed with *amtA M1* and *amtA M2.* Growth phenotype of these two transformants was observed using the spotting assay (Figure 1D) and by streaking for single colonies on low ammonium media (Figure S3). It is clear that the triple deletion strain transformed with these mutants grow on low ammonium media. Wild type *amtA* (henceforth referred to as *amtA WT*), as well as *amtA M1,* and *amtA M2* were integrated at the *MEP2* locus such that the expression is driven by *MEP2* promoter. The ability of these strains to grow on low ammonium media was tested by spotting assay (Figure 1E). These results are similar to the results obtained using single copy plasmid transformants.

We then carried out growth kinetics of these strains when grown in minimal medium containing 0.1% proline as the sole nitrogen source and on low ammonium media (Figure 1F, 1G). The growth assay in minimal medium containing 0.1% proline as the sole nitrogen source was monitored at an interval of 1 hr for a period of 36 hrs (Figure 1F). All the strains show a similar growth pattern with a growth rate of 0.11 hr^−1^. For the growth assay in a low ammonium media (Figure 1G), the experiment was carried out for 48 hrs. The triple deletion strain and the strain expressing *amtA WT* did not show any growth. The strain expressing *MEP2* grew with a growth rate of 0.03 hr^−1^. In comparison, strains expressing *amtA M1* and *amtA M2* grew with a growth rate of 0.1 hr^−1^ and 0.08 hr^−1^, respectively. However, strain expressing *MEP2* showed a growth lag of 12 hrs whereas the strains with *amtA M1* and *M2* showed a growth lag of 20 hrs. Further, the final biomass of strain expressing *MEP2* was much lower than that of strains expressing *amtA M1* and *amtA M2* at saturation. The observed difference in growth profile between *MEP2* and *amtA* mutants, clearly reflects a difference in their functionality as ammonium transporters. However, the underlying cause for the observed difference is difficult to decipher.

We also determined the ability of strains expressing *amtA WT*, *amtA M1*, *amtA M2*, and *MEP2* to take up ammonium from the medium as a function of time (Figure 1H). Strains were grown in low ammonium media, and the ammonium remaining in the medium as a function of time was estimated. While triple deletion strain or the strain expressing *amtA WT* failed to take up ammonium (Figure 1H), *amtA M1* and *amtA M2* strains showed ammonium uptake activity comparable to *MEP2* strain. At the end of 3 hours, the ammonium concentration is close to 0 in the medium in which strains expressing *amtA M1*, *amtA M2*, *MEP1,* and *MEP2* were grown. Whereas the external ammonium concentration remained close to the initial concentration in the medium in which triple deletion strain and strain expressing *amtA WT* were grown. This observation is in concordance with the growth phenotype on low ammonium where only strains expressing *amtA M1*, *amtA M2*, *MEP1*, and *MEP2* grew.

### Both synonymous and nonsynonymous substitutions are necessary to confer functionality to *amtA M1* and *amtA M2*

Comparison of nucleotide sequences of *amtA WT*, *amtA M1* and *amtA M2* indicated that *amtA M1* has 3 nonsynonymous (Q89L, A134V, S311T) and 2 synonymous (I301I, S350S) mutations and *amtA M2* had two nonsynonymous (P129T, V334I) and one synonymous (K288K) mutations. 31 and 7 different combinations of these single mutations are possible in *amtA M1* and *amtA M2,* respectively. By site directed mutagenesis, plasmid clones bearing these combinations of nonsynonymous and synonymous substitutions of *amtA M1* and *amtA M2* respectively, were generated. The functionality of these mutant clones was tested by carrying out the growth kinetics by growing the transformants on low ammonium media, as described in the previous section (Figure 2). Plasmid clones of *amtA M1* bearing individual nonsynonymous or synonymous substitutions did not grow on low ammonium media. Nonsynonymous substitutions either in pairs of two or three also did not grow (Figure 2A). Similarly, synonymous mutations either alone or both together did not confer activity. Two nonsynonymous mutations and one synonymous mutation in all the combinations did not confer functionality. Three nonsynonymous substitutions with any one of the synonymous substitutions also did not show the activity. The functionality was observed only when all the three nonsynonymous and two synonymous substitutions are present together (Figure 2A). The above analysis indicates that both synonymous and nonsynonymous substitutions are necessary to confer functionality to *amtA M1*.

**Figure 2:**
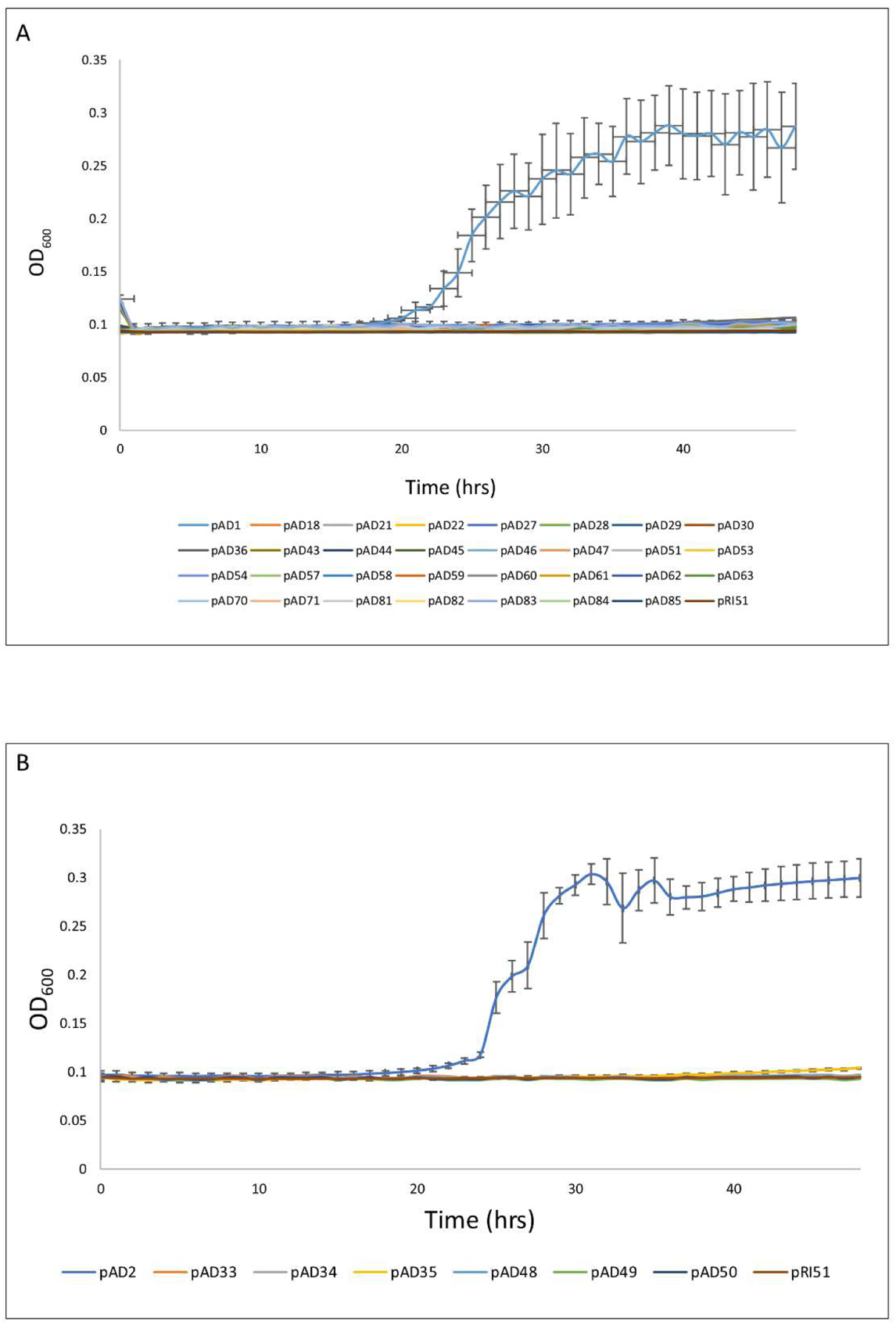
Combination of synonymous and nonsynonymous mutations is essential for *amtA M1* and *amtA M2* functionality. **(A) Growth kinetics of transformants on low ammonium.** Different transformants pre-grown in 0.1% proline medium, were grown in low ammonium media, and OD_600_ was measured as a function of time for 48 hours. Each line represents a transformant bearing a plasmid clone with specific substitution mutation(s) in *amtA WT* corresponding to *amtA M1*. The resulting substitutions are: pAD1: *amtA_Q89L, A134V, I301I, S311T, S350S_*_;_ pAD18: *amtA_Q89L_*_;_ pAD21: *amtA_A134V_*_;_ pAD22: *amtA_S311T_*_;_ pAD27: *amtA_Q89L, S350S_*_;_ pAD28: *amtA_Q89L, A134V_*_;_ pAD29: *amtA_A134V, S311T_*_;_ pAD30: *amtA_Q89L, S311T_*_;_ pAD36: *amtA_Q89L, A134V, S311T_*_;_ pAD43: *amtA_A134V, I301I_*_;_ pAD44: *amtA_I301I_*_;_ pAD45: *amtA_Q89L, A134V, I301I, S311T_*_;_ pAD46: *amtA_S350S_*_;_ pAD47: *amtA_Q89L, A134V, S311T, S350S_*_;_ pAD51: *amtA_I301I, S350S_* pAD53: *amtA_A134V, S350S_*_;_ pAD54: *amtA_Q89L, A134V, I301I, S311T, S350S_*_;_ pAD57: *amtA_S311T, S350S_*_;_ pAD58: *amtA_A134V, I301I, S311T_*_;_ pAD59: *amtA_I301I, S311T_*_;_ pAD60: *amtA_I301I, S311T, S350S_*_;_ pAD61: *amtA_Q89L, I301I, S350S_*_;_ pAD62: *amtA_A134V, I301I, S350S_*_;_ pAD63: *amtA_A134V, S311T, S350S_*_;_ pAD70: *amtA_Q89L, A134V, S350S_*_;_ pAD71: *amtA_Q89L, A134V, I301I_*_;_ pAD81: *amtA_Q89L, I301I, S311T_*_;_ pAD82: *amtA_Q89L, S311T, S350S_*_;_ pAD83: *amtA_A134V, I301I, S311T, S350S_*_;_ pAD84: *amtA_Q89L, I301I, S311T, S350S_*_;_ pAD85: *amtA_Q89L, A134V, I301I, S350S_*_;_ pRI51: *amtA* **(B) Growth kinetics of transformants on low ammonium.** Different transformants pre-grown in 0.1% proline medium, were grown in low ammonium media, and OD_600_ was measured as a function of time for 48 hours. Each line represents a transformant bearing a plasmid clone with specific substitution mutation(s) in *amtA WT* corresponding to *amtA M2*. The resulting substitutions are: pAD2: *amtA_P129T, K288K, V334I_*_;_ pAD33: *amtA_P129T_*_;_ pAD34: *amtA_V334I_*_;_ pAD35: *amtA_P129T, V334I_*_;_ pAD48: *amtA_K288K_*_;_ pAD49: *amtA_P129T, K288K_*_;_ pAD50: *amtA_K288K, V334I_*

It was observed that plasmid clones bearing either nonsynonymous or synonymous mutations of *amtA M2* individually could not grow on low ammonium (Figure 2B). Combination of any of the two mutations, synonymous or nonsynonymous did not confer functionality. The functionality was observed only when all the 2 nonsynonymous and 1 synonymous mutations were present (Figure 2B). These observations clearly demonstrate that nonsynonymous or synonymous mutation in isolation do not confer the functionality and the combination of both are necessary for the function of *amtA M2* as well as *amtA M1*. To the best of our knowledge, this is the first report where a gain of function is dependent on both types of substitutions.

### AmtA wild type and its mutant derivatives exhibit differential behaviour in plasma membrane and sensitivity to proteolytic cleavage

The membrane localization pattern of AmtA M1 and AmtA M2 was studied by using strains with yEGFP tagged to the C-terminus of respective proteins. The functionality of the mutants after yEGFP tagging was confirmed by testing their ability to grow on low ammonium media (Figure S4). Fusing yEGFP to the C-terminus of AmtA mutant derivatives did not hamper their biological function (Figure S4). GFP fluorescence signal was detected in the plasma membrane of *amtA M1::GFP* and *amtA M2::GFP* strain (Figure 3A). GFP signal was also observed in the subcellular compartment in the strain expressing *amtA M2::GFP* similar to what is observed for the strain expressing *amtA WT::GFP* (Figure 3A). Subcellular localization of human ammonium transporters RhAG and RhCG in *S. cerevisiae* has been reported elsewhere [72]. As of now, it is not clear if the subcellular GFP signal observed in cells expressing *amtA WT*, and *amtA M2* is a degraded product or the fusion protein.

**Figure 3:**
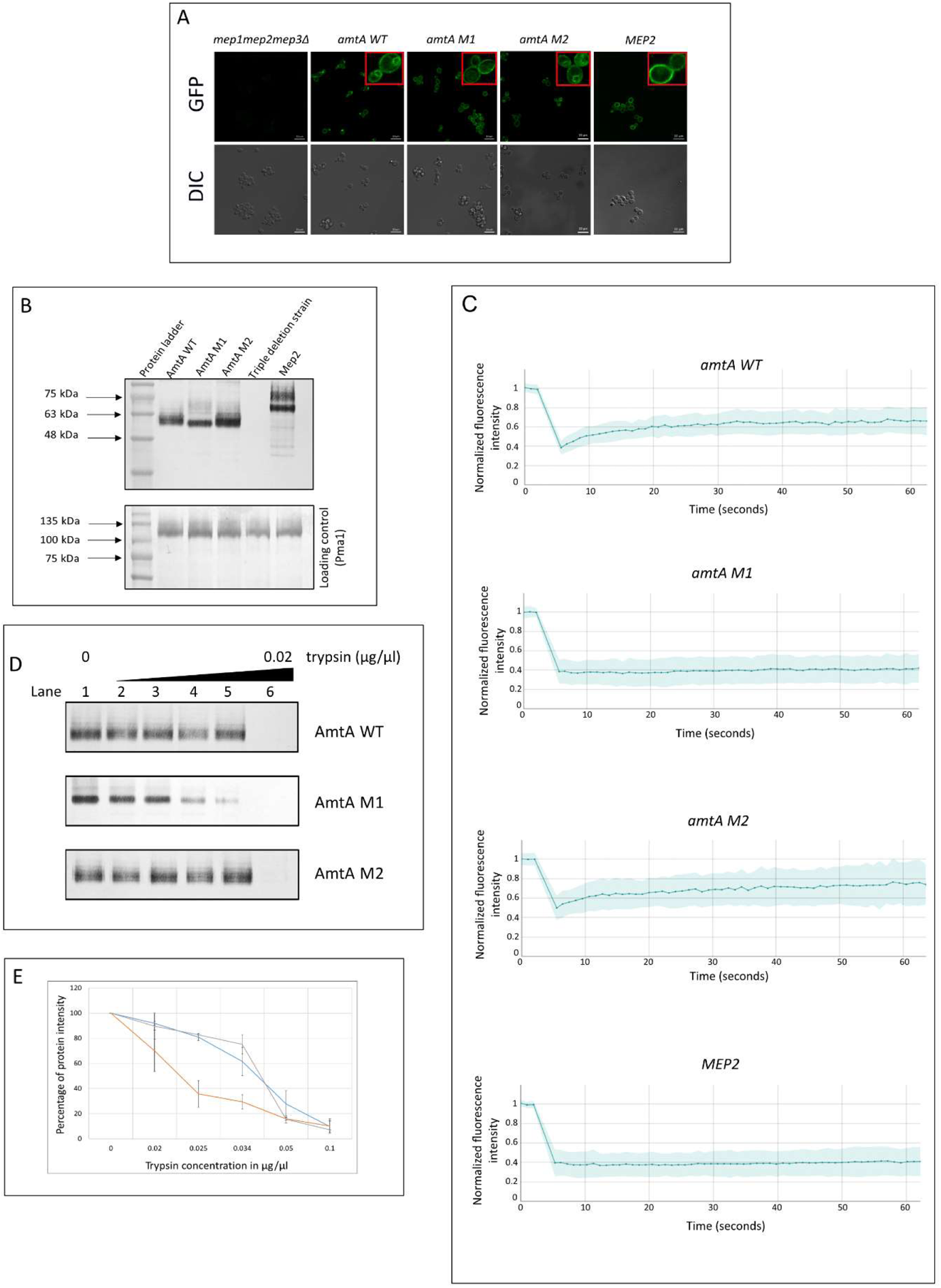
*amtA WT*, *amtA M1* and *amtA M2* differ in their proteolytic sensitivity and plasma membrane mobility. **(A) Expression and localization of yEGFP-tagged proteins.** Strains with yEGFP tagged proteins were grown in 0.1% proline media for 22 hours. The localization of the yEGFP tagged proteins was visualized using live fluorescence microscopy. The top panel represents the GFP fluorescence images, and the bottom panel represents DIC images. **(B) Protein expression in the membrane fraction.** The membrane fraction was isolated from strains grown in 0.1% proline media for 22 hours, and western blotting was performed. Anti-GFP antibody was used to detect the GFP tagged protein. Pma1 antibody was used to detect Pma1 protein that served as a loading control. (C) Fluorescent Recovery After Photobleaching (FRAP). *yEGFP*-tagged *amtA WT*, *amtA M1*, *amtA M2*, and *MEP2* strains were subjected to FRAP for 60 seconds. The fluorescence intensity at each data point is normalized with respect to the fluorescence intensity at photobleaching. **(D) Digestion pattern after trypsin digestion of membrane fraction.** Membrane fractions of *amtA WT*, *amtA M1*, *amtA M2*, and *MEP2* strains were subjected to trypsin digestion. No trypsin is used in lane 1, and increasing concentration of trypsin, 0.004 µg/µl, 0.005 µg/µl, 0.0067 µg/µl, 0.01 µg/µl, and 0.02 µg/µl, was used from 2nd lane onwards. **(E) Quantitative analysis of protease digestion pattern after trypsin digestion of membrane fraction.** Membrane fractions of *amtA WT*, *amtA M1*, *amtA M2*, and *MEP2* strains were subjected to trypsin digestion. No trypsin is used in lane 1, and increasing concentration of trypsin0.004 µg/µl, 0.005 µg/µl, 0.0067 µg/µl, 0.01 µg/µl, and 0.02 µg/µl, was used from 2nd lane onwards. The percentage of protein intensity with respect to the protein intensity present without trypsin is plotted against trypsin concentration.

To corroborate the observations that AmtA and its derivatives are localized in the plasma membrane, we carried out a subcellular isolation of plasma membrane from strains grown in 0.1% proline, followed by western blotting using antibodies against GFP (Figure 3B). It is pertinent to mention that membrane fraction was treated with SDS and 8M urea and incubated at room temperature instead of heating at 100°C before subjecting it to SDS electrophoresis, to avoid protein aggregation [73]. Plasma membrane ATPase 1 (Pma1) protein was used as a loading control. It can be observed that AmtA WT, AmtA M1, and AmtA M2 are present in the membrane fraction (Figure 3B). The normalized intensity of AmtA WT, AmtA M1, and AmtA M2 with respect to Pma1 was found to be 1.5, 1.6, and 2.1, respectively. The expected molecular weight of the AmtA fusion protein is 77 kDa. The observed molecular weight obtained from the above data is 62 kDa. It is commonly observed that membrane proteins exhibit anomalous mobility on SDS PAGE to the extent of approximately 30% [74].

It has been previously reported that synonymous codons can result in the synthesis of conformationally heterogeneous protein products [75]. This can result in a difference in disposition between AmtA M1 and AmtA M2. For example, ammonium transporters are known to exist as trimers and a difference in the conformation can affect the trimerization [76][77]. It is also possible that the lipid-protein interaction in the plasma membrane could have a deleterious effect on the functionality because of a difference in the disposition of AmtA M1 and AmtA M2 in the plasma membrane. We decided to probe the above possibilities by carrying out analysis to determine if there are any difference in the conformation of the proteins and by extension a difference in the lateral diffusion of the proteins in the plasma membrane. Fluorescence Recovery After Photobleaching (FRAP) analysis was performed to determine whether there exists a difference in lateral diffusion between AmtA WT, AmtA M1 and AmtA M2 in plasma membrane, a milieu in which it is functional. The pattern of recovery after photobleaching was determined by measuring the normalized fluorescence intensity as a function of time. The measurement was done for a total of 60 seconds on 37 samples for each strain (Figure 3C). We observed that AmtA M1 and Mep2 fail to recover after photobleaching, with a recovery rate of 4.76% and 6.54% respectively, suggesting that their mobility is restricted. On the other hand, AmtA WT recovered to the extent of 41.77% while AmtA M2 recovered to the extent of 48.82%. This suggests that both AmtA WT and AmtA M2 have a mobile fraction. These results suggest that functionality is not directly correlated with recovery rate. One way to reconcile this apparent dichotomy is by assuming that they differ in their conformation.

To further probe the differences between AmtA WT, AmtA M1, and AmtA M2, we looked at the sensitivity of these proteins to trypsin digestion. If there are conformational differences between AmtA WT and mutants, the kinetics of digestion in the presence of a varying concentration of trypsin is expected to vary. This kinetics was monitored by probing the protein using GFP antibody on Western Blot, as GFP does not get digested by trypsin [78][79]. Thus, if any, the difference in the kinetics of trypsin digestion ought to be due to a difference in the conformation of the proteins. Figure 3D shows a representative result of the experiment. An average of four experiments are represented in Figure 3E. It can be observed that the amount of trypsin needed to digest 50% of the protein is twice for AmtA WT and M2 compared to M1.

This data reveals that AmtA M1 is more susceptible to trypsin as compared to AmtA WT and AmtA M2, suggesting a difference in the conformation between AmtA WT and its derivatives. The difference in recovery rate after photobleaching and the sensitivity to trypsin digestion of AmtA M1 from that of AmtA WT and AmtA M2 suggests that the conformation of AmtA M1 is different from that of the AmtA WT and AmtA M2. On the other hand, we could not detect a discernible conformational difference between AmtA WT and AmtA M2 using the above approaches.

The role of synonymous mutations mainly manifests at two different levels. First, at the level of protein synthesized, which normally is a function of mRNA stability or structure because of synonymous mutations. We calculated the free energy change of the *WT* and the *mutants’* mRNA using the online software RNA structure Fold [80]. The ΔG values for *amtA WT, amtA M1,* and *amtA M2* are −338.1, −340, and −339.7, respectively. This change is unlikely to cause any significant alteration in the stability of the mRNA. Further, the concentration of protein localized on the membrane is comparable to that of Mep2. These two observations seem to rule out the above possibility. The second level at which synonymous mutations alter protein functionality is by changing translational kinetics, which affects co-translational folding, thereby causing a conformational change. Given that nonsynonymous change is also required for functionality, we are inclined to suggest that synonymous mutations’ role is to introduce a conformational change. There is sufficient literature wherein the above mechanisms have been studied [81][82][83][75].

### AmtA M1 and AmtA M2 show different sensitivity to methylamine toxicity and distinct mechanisms of ammonium transport

AmtA M1 and M2 show ammonium uptake activity and biochemical and microscopic examination suggest conformational differences. How does one reconcile these differences? It is possible that the difference in the conformation is a reflection of their biological activity. For example, *S. cerevisaie* Mep proteins take up methylamine, albeit to different extent [84][85]. It was also shown that strain expressing only *MEP1* but not *MEP2* alone or *MEP3* alone, confer sensitivity to methylamine when cells are grown in 0.1% proline as the sole source of nitrogen [86][84] (Figure 4A). We have confirmed this observation (Figure 4A). The inability of Mep2 to confer methylamine toxicity is attributed to the conserved H194 which is present in Mep2 but not in Mep1, which has glutamate at the corresponding position instead of histidine [84]. Accordingly, the ability of Mep1 to cause methylamine toxicity is abolished by replacing E194 by histidine. Substituting glutamate instead of histidine at position 194 in Mep2 confers the methylamine toxicity. These studies point out that glutamate at position 194 corresponding to the conserved histidine is crucial for imparting the methylamine toxicity [84].

**Figure 4:**
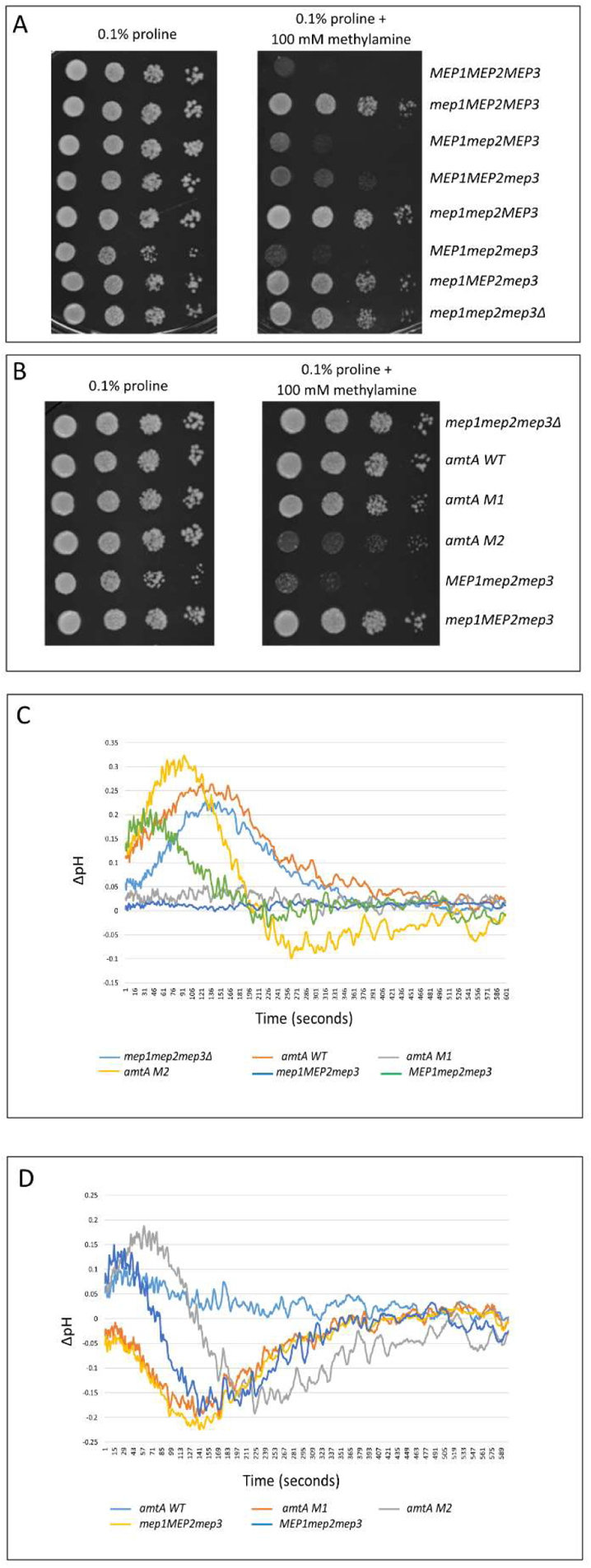
AmtA M1 and AmtA M2 differ in their ammonium transport mechanism and methylamine sensitivity. **(A) Growth of strains on methylamine.** 5 µl of the strains pre-grown on 0.1% proline media to a cell density of 10^7^/ml were serially diluted and were spotted onto 0.1% proline media (left panel), and synthetic minimal medium with 0.1% proline and 100 mM methylamine (right panel). The growth pattern was photographed after 4 days of incubation at 30°C. **(B) Growth of strains on methylamine.** 5 µl of the strains pre-grown on 0.1% proline media to a cell density of 10^7^/ml were serially diluted and were spotted onto 0.1% proline media (left panel), and synthetic minimal medium with 0.1% proline and 100 mM methylamine (right panel). The growth pattern was photographed after 4 days of incubation at 30°C. **(C) Intracellular change in pH after ammonium addition as a function of time.** ΔpH was measured using pH-sensitive GFP as a function of time and plotted for different strains. ΔpH = (Intracellular pH of strain when exposed to ammonium - intracellular pH of strain when exposed to water). **(D) Intracellular change in pH after ammonium addition as a function of time.** ΔpH was measured using pH-sensitive GFP as a function of time and plotted for different strains. ΔpH = (Intracellular pH of strain when exposed to ammonium - intracellular pH of strain when exposed to water) - (Intracellular pH of triple deletion strain when exposed to ammonium - intracellular pH of triple deletion strain when exposed to water).

AmtA WT, AmtA M1, and AmtA M2 have histidine at the conserved position 194. Based on the above, AmtA WT and its mutant derivatives are not expected to confer methylamine toxicity to triple deletion strain. To investigate whether the above expectation holds good, we carried out a similar analysis with triple deletion expressing *amtA WT*, *amtA M1*, or *amtA M2* individually. As a positive and negative control, we included triple deletion strain expressing *MEP1* or *MEP2* individually (Figure 4B). It is observed that AmtA M2 confers methylamine toxicity while AmtA WT and AmtA M1 do not confer methylamine toxicity. These observations suggest that the glutamate at position 194 *per se* is not necessary for conferring methylamine toxicity. Even if histidine is present at this position, it can confer methylamine toxicity.

Since AmtA WT, AmtA M1 and AmtA M2 show different response to methylamine toxicity, we were curious to see whether AmtA M1 and AmtA M2 behave differently with respect to the mechanism of ammonium transport. We used a ratiometric GFP, which is sensitive to pH, to measure the cytosolic pH to monitor the intracellular pH as function of exposure of cells to ammonium pulse at pH 6.1 as demonstrated in a recent study [87]. In this study intracellular pH as a function of ammonium transport was reported using pHluorin as a probe in *S. cerevisiae* [87]. Yeast cells grown at pH 6.1 were used to measure the effect of ammonium transporter Mep1, Mep2, and Mep2 mutants on the cytosolic pH of the cells, after the addition of ammonium at a concentration of 2 mM. We followed a similar protocol where cells were grown in a medium of pH 6.1 containing 0.1% proline as the sole nitrogen source. At OD_600_ 1.0, ammonium was added to a concentration of 2 mM, and the intracellular ratiometric fluorescence was recorded as a function of time.

A triple deletion strain served as a negative control. Figure 4C shows the profile of all the strains after subtracting the values obtained from the control (addition of water instead of ammonium). This reflects the change in the intracellular pH only due to the presence of ammonium in the medium. Triple deletion strain and *amtA WT* strain respond similarly. Strains expressing *amtA M1* and *MEP2* behave identically, while the pattern observed of *amtA M2* and *MEP1* strains are similar, although they are different in magnitude. Figure 4D provides the profile of intracellular pH after subtracting the values obtained from the triple deletion strain. In effect, this would get rid of any change in the pH profile caused by any phenomena other than ammonium transport. Here, the absolute profile has changed, but the pattern between the strains is the same. It can be observed that *amtA M1* and *MEP2* strains respond identically. This is not surprising as they both transport ammonium and are not sensitive to methylamine. On the other hand, strains expressing *amtA M2* and *MEP1* transport ammonium and are sensitive to methylamine, and their intracellular pH response to ammonium addition is similar. Thus, we have been able to demonstrate functional equivalence between AmtA M1 and Mep2 as well as between AmtA M2 and Mep1, even with respect to intracellular pH profile.

The mechanisms of translocation of ammonium are measured either using homologous system or by heterologous system. In a heterologous system, the transporters are expressed routinely in Xenopus oocytes, and electrophysiological experiments are performed. A second approach has been to reconstitute purified ammonium transporters into liposomes and then follow the translocation [88][89][90][87]. In either case, current is measured at a fixed voltage as a function of ammonium translocation. It has been observed that the ammonium translocation causes a change in the current as early as few seconds, and thereafter it reaches a steady state. This pattern was also observed when intracellular pH was monitored as a function of time in response to ammonium addition. Using both the approaches, it was inferred that Mep1 and Mep2 translocate ammonium through electrogenic and neutral mechanisms, respectively [87]. Based on the above and the results presented here, we propose that AmtA M1 translocates ammonium through a neutral mechanism while AmtA M2 translocates ammonium through an electrogenic mechanism.

## Discussion

Deciphering the role of epistasis between nonsynonymous mutations has been the mainstay of understanding structure-function relationship and evolution of proteins [91]. Because of technological advancement, it is now possible to study higher order epistatic interaction involving more than two amino acid residues [92]. However, it has now become increasingly clear that even synonymous substitutions can result in altered conformation leading to functional alteration [75], thereby making it difficult to study protein evolution. Thus far, there is no direct experimental evidence in support of an epistatic interaction between synonymous and nonsynonymous substitutions leading to functional alteration. Based on the results presented here, it is clear that protein evolution can’t be understood by studying the consequence of nonsynonymous and synonymous substitutions in isolation. It was suggested that to understand the role of epistasis in proteins evolution, the measurements have to be made in the native host [93]. However, we contend that since most of the studies on epistasis are performed in native host or host which are permissive for the functional expression of proteins, the interaction between synonymous and nonsynonymous substitutions that otherwise exists, would have escaped detection. For example, as discussed in the introduction, triple deletion strain serves as a permissive host for the functional expression of ammonium transporters. That is, studying the role of epistasis in the evolution of AmtA in *D. discoideum* would have not yielded the results presented here, if yeast were to be a permissive host of *D. discoideum* AmtA. The biological meaning of such context dependent behaviour of synonymous mutations is highlighted in a study wherein the authors demonstrated that synonymous substitution can function in a tissue specific manner [83]. Based on the above, we suggest that complementation across incompatible genomes can be a powerful approach to decipher the trajectory of protein evolution.

A possible mechanism of epistasis between synonymous and nonsynonymous mutation observed in this study is schematically represented in Figure 5. For the ease of understanding, epistasis in *amtA M2*, is taken as an example. We conjecture that a change from Lysine codon AAA to AAG has resulted in an increase in translational speed. This is based on the observation that in *S. cerevisiae*, the number of tRNA genes corresponding to AAA is 14 as compared to 7 for AAG [94]. Green, red and blue spheres represent hypothetical amino acid residues. In the functional protein, green interacts with red. Because of the increased translational speed, green interacts with blue instead of red leading to a non-functional protein. This nonspecific interaction is suppressed and the specific interaction between green and red is restored because of P129T and V334I substitutions. Unlike synonymous mutations, nonsynonymous substitutions can alter the folding dynamics either by changing the translation speed or through residue-residue interaction or both. Because a direct interaction between Threonine and Isoleucine is not possible, we suspect that their effect is mediated through an as yet unknown mechanism.

**Figure 5:**
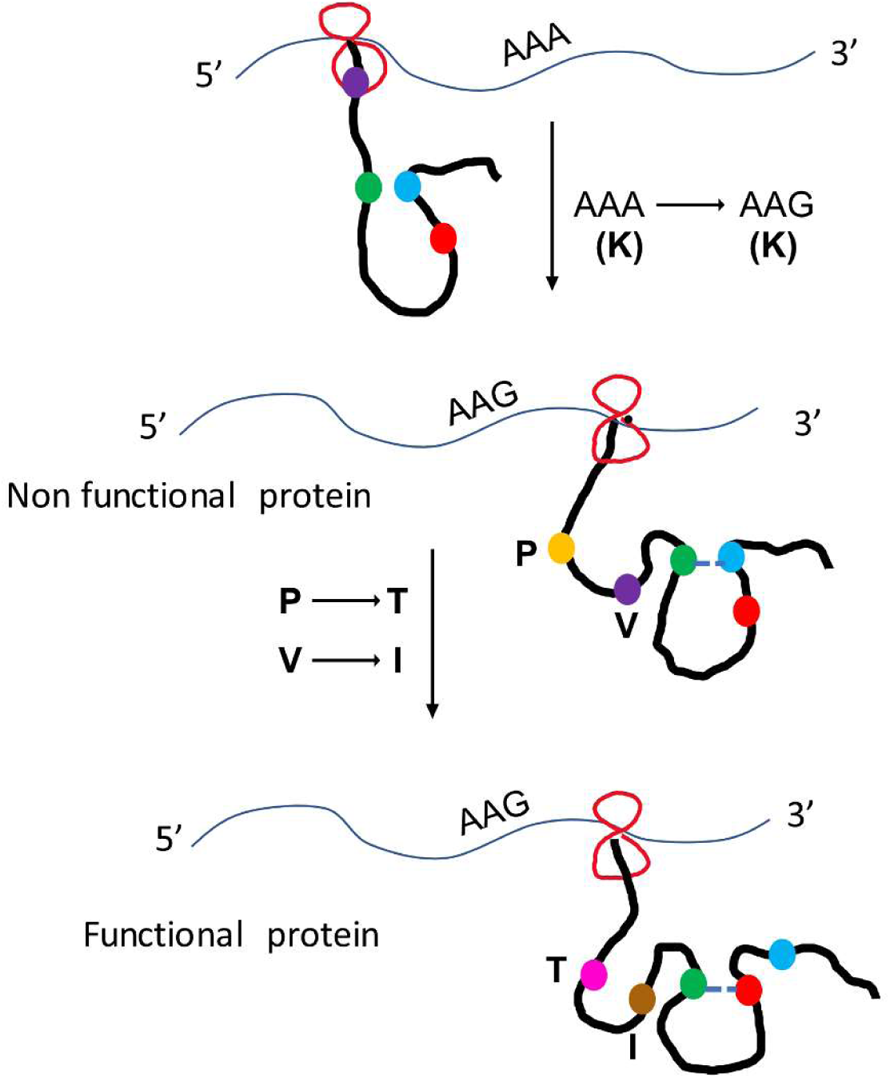
Representative mechanism of AmtA M2 gain of function in *S. cerevisiae*.

Once a genome acquires a new gene through processes such as genome duplication, horizontal gene transfer, the gene can be modified in very many ways to fulfil the demands of selection. Of these modifications, the role of single nucleotide mutation leading to nonsynonymous substitutions is the mainstay of protein evolution [91]. This is because protein evolution is contingent upon the interaction between residues in the same protein (intragenic) or on residues of the interacting proteins (intergenic). During the early days of molecular evolutionary studies, it was proposed that wide range of amino acids can be substituted without altering the functionality only at some sites, but not all sites [95]. That is, evolution of different sites in a protein is constrained to different extent, because of the co-operative nature of interaction between any pairs of amino acids, a phenomenon commonly termed as intragenic epistasis, first experimentally observed in tryptophan synthase [96]. In fact, more often, the interaction can be between three or more number of amino acids, a phenomenon termed as higher order epistasis, observed first in Lysozyme [97]. The study of pairwise or higher order epistasis in protein evolution is intensely investigated in the past two decades to understand protein evolution with a larger goal of mapping genotype to phenotype [92].

It is now well documented that synonymous substitutions are not neutral and play a significant role in protein biogenesis by altering translational fidelity, translational rate, co translational folding and therefore can eventually have profound effect on organismal fitness [81][82][83][75][98]. Recent studies have also demonstrated that intragenic epistasis between synonymous mutations can constrain the evolutionary trajectory[99][100][101]. Surprisingly, in one of the above studies, the mechanistic basis of the increase in fitness caused by the synonymous and nonsynonymous mutations could not be discriminated [99]. Much of our knowledge on the biological consequences of synonymous substitution comes from studies where the effect of synonymous codons is determined while keeping the rest of the amino acid sequence context intact. But this rarely happens during protein evolution. In a real scenario, protein suffers from many mutational events, following which, functionally important amino acid residues are selected. While the substitutions of amino acids in orthologues can be rationalised using mutational (deep mutational scanning, ancestral protein resurrection) and biophysical techniques, the underlying chemical logic for the conservation of codons in the context of varying sequences is hard to decipher [91]. That is, in the absence of a priori theoretical understanding, the predictive power to empirically determine the effect of synonymous substitution on fitness, under the conditions of varying sequence context is lacking. A case in point is the observation that a synonymous substitution of I507 from ATC to ATT altered drug sensitivity of an otherwise non-functional ΔF508 CFTR allele [102].

More recently, synonymous SNPs have been strongly correlated with many genetic diseases, but the underlying mechanisms have not been clearly elucidated. It has been proposed that synonymous mutations act as driver mutations, implying that the disease-causing nonsynonymous mutations can arise in the background of synonymous mutations [103][104]. In the light of what we have observed, it is now possible to look for specific nonsynonymous mutations in the background of driver mutations. Obtaining in depth understanding of epistasis between synonymous and nonsynonymous substitutions and how these interactions influence protein folding is the key in unearthing the trajectory of molecular evolution and eventually phenotypic evolution.

The study presented here goes far beyond just addressing the canonical problem of protein evolution. Understanding intragenic epistasis alone may be insufficient to address the evolutionary trajectory of a protein following such events as gene duplication and more so HGT. This is because, as mentioned in the introduction, evolution following gene duplication occurs within the related genome (Darwinian evolution) as opposed to HGT where evolution occurs in an alien genome (non-Darwinian evolution). Based on theoretical analysis, it has been proposed that HGT played a crucial role in the evolution of universality and optimality, the defining features of genetic code [105]. In this context, our study has raised more questions than has answered. For, example, would Mep2 be functional in *D. discoideum*? Would it be possible at all to obtain a functional AmtA only by introducing nonsynonymous mutations, without the aid of synonymous mutations? Can bifunctionality (ammonium and methylamine transport) be introduced by altering translation kinetics? Answer to such questions would not only provide insights into the possible early evolutionary events but also may provide insights into the bottlenecks often encountered in obtaining functionally active proteins using heterologous expression systems. The experimental system we have studied here is amenable to experimental analysis and therefore can be fine-tuned to address biological problems of both academic and applied interest.

## Materials and Methods

### Media Components and Growth conditions

The yeast cultures were grown in YPD, and appropriate Synthetic Drop out Media. The bacterial (*E. coli*) cultures were grown in Luria Broth Agar and liquid media and complemented with Ampicillin at a concentration of 75 µg/ml whenever required. Yeast Extract, LB Broth, and Agar were purchased from Hi-Media Laboratories. Bactopeptone and Yeast Nitrogen Base were obtained from Difco Detroit, Michigan.

#### Yeast Extract Peptone Dextrose (YPD) medium

Yeast extract - 0.5%, Peptone - 1%, Dextrose - 2%, Agar – 2%

#### Synthetic complete and drop out glucose medium

Glucose - 2%, Ammonium sulfate +Yeast Nitrogen Base (YNB) without amino acids mix (3:1) - 0.66 gm/100 ml, Amino acid mix - 0.05 gm/100 ml, Agar - 2%

#### Amino acid mix

Adenine - 0.8 gm, Uracil - 0.8 gm, Tryptophan - 0.8 gm, Histidine - 0.8 gm, Arginine - 0.8 gm, Methionine - 0.8 gm, Leucine - 1.2 gm, Tyrosine - 1.2 gm, Lysine - 1.2 gm, Phenyl alanine - 2.0 gm, Threonine - 8.0 gm, Aspartic Acid - 4.0 gm, Valine - 1.5 gm, Isoleucine - 0.3 gm Complete amino acid mixture contains all the above-mentioned amino acids and nitrogen bases. A particular drop out amino acid mixture lacks that specific component.

#### Minimal media with 500 µM Ammonium

Ammonium Sulfate – 500 µM, Yeast Nitrogen Base (without ammonium sulfate) – 0.17%, Glucose – 2%, Agar – 2%

For media containing different ammonium concentrations, only ammonium sulfate concentration varies accordingly.

### 0.1% Proline media

Proline – 0.1%, Yeast Nitrogen Base (without ammonium sulfate) – 0.17%, Glucose – 2%, Agar – 2%

### 0.1% Proline 200 mM methylamine medium

Proline – 0.1%, Methylamine hydrochloride – 200 mM, Yeast Nitrogen Base (without ammonium sulfate) – 0.17%, Glucose – 2%, Agar – 2%

For media containing different methylamine concentrations, only methylamine hydrochloride concentration varies accordingly.

### DNA modification, electrophoresis and Western blotting

Tris-EDTA, Amino Acids, and the reagents used in Agarose Gel Electrophoresis were obtained from Sigma-Aldrich USA. The restriction enzymes were obtained from Fermentas, and DNA polymerases were obtained from NEB; all other chemicals used were of analytical grade. Pma1 was probed with Mouse Pma1 Monoclonal Antibody (40B7), yEGFP was probed using Anti-GFP, from mouse (Roche #11814460001) antibody and Anti-Mouse IgG −Alkaline Phosphatase antibody (A9316) was used as the secondary antibody.

### Plasmid Isolation from *E. coli*

The plasmid was isolated from the *E. coli* by the Alkaline Lysis Method given by [106].

### Yeast Transformation

A single colony was inoculated into 5 ml YPD and incubated at 30°C with shaking till O.D. at 600 nm reaches up to 0.5. Cells from 1.5 ml of the culture were harvested by centrifugation at 13000 rpm for 1 minute. The cells were washed thrice with 200 μl of TE buffer (10 mM Tris, 1 mM EDTA pH 8.0). Then the cells were washed two times with 200 μl of 100 mM lithium acetate. The cells were re-suspended in 150 μl of 100 mM lithium acetate, and DNA was added to these cells along with 5 μl of single-stranded DNA. The cultures were incubated at 30°C for 5 minutes. To this mixture, 1 ml of 40% PEG (3350) was added, and the tubes were incubated at 30°C for 1 hr. After 1 hr of incubation, cells were subjected to heat shock at 42°C for 20 minutes. Cells were harvested by centrifugation at 13,000 rpm for 5 minutes and washed twice with TE buffer. The cells were then resuspended in 100 μl TE buffer and plated onto the appropriate medium.

### Yeast genomic DNA isolation

A single colony of yeast was inoculated into 5 ml of YPD broth and grown at 30°C for approximately 20 hours on an orbital shaker. Cells from this culture were pelleted in a microcentrifuge tube. The pellet was resuspended in 200 μl of lysis buffer (2% Triton X-100, 1% SDS, 100 mM NaCl, 10 mM Tris-HCl pH 8.0, and 1 mM EDTA pH 8.0). 0.3 gm of acid-washed glass beads and 200 μl of phenol-chloroform-isoamyl alcohol mix were added to the above tubes. The tubes were then vortexed for 3-5 minutes in intervals of 30 seconds and intermittent incubation on ice. After adding 200 μl of TE buffer, the tubes were centrifuged for 15 min at 13,000 rpm at 4°C. The aqueous layer was transferred to a fresh tube, and DNA was precipitated by adding 1 ml of absolute ice-cold ethanol. The samples were allowed to precipitate for 5 min at 4°C and then centrifuged for 10 min at 13,000 rpm at 4°C. DNA pellets were air-dried and re-suspended in 400 μl TE buffer with 3 μl RNAse (10 mg/ml) and kept for 5 minutes at 37°C. After this, the DNA was reprecipitated by adding 10 μl of 4 M ammonium acetate and 1 ml of absolute ice-cold ethanol. After centrifugation at 4°C for 10 minutes at 13,000 rpm, the DNA pellet was dissolved in 25 μl TE buffer.

### Error prone PCR

The PCR condition for *amtA* amplification comprised of initial denaturation at 95°C for 7 minutes, followed by 30 cycles of denaturation at 95°C for 1 minute, annealing at 55°C for 1 minute and extension at 72°C for 3 minutes. Final extension was carried out at 72°C for 14 minutes. In order to introduce random mutations, dTTP and dCTP were used at a concentration of 4mM while dATP and dGTP at a concentration of 2 mM. Also, MnCl_2_ was added to the PCR mix. Error Prone PCR was carried out using different concentrations of MnCl_2_ to determine the optimal concentration of MnCl_2_. It was observed that the yield of PCR product was maximum at a concentration of 0.35 mM and 0.45 mM of MnCl_2_. Further PCR experiments for mutagenesis were carried out at 0.35 and 0.45 mM MnCl_2_.

### Disruption/Integration of ORF

Specific ORF was disrupted/integrated in the strain using KanMX4 cassette scoring for Geneticin resistance or using hphMX6 cassette scoring for Hygromycin resistance. The primers were used to amplify the KanMX4 cassette from the pUG6 plasmid or the hphMX6 cassette from the pUG75 plasmid. The transformants were selected onto a YPD plate containing geneticin antibiotic or hygromycin antibiotic. The putative transformants were tested for the gene disruption/integration by diagnostic PCR using an internal primer of KanMX4 cassette or hphMX6 cassette and also using the primers that give a product if ORF is disrupted/integrated, while the wild type strain will not give any PCR product.

### Ammonium estimation assay

The strains were grown in media containing 0.1% proline as the sole nitrogen source until the OD_600_ reaches 1.0. The cells were collected and washed twice with sterile distilled water and transferred to media containing 500 µM ammonium and 0.1% proline as the nitrogen source. The external ammonium concentration was estimated as a function of time for 3 hours using the below-mentioned principle (Ammonia Assay Kit - Sigma-Aldrich). At each time point, cells were collected, filtered and the filtrate was taken for estimating ammonium. Ammonia reacts with α-ketoglutarate and reduced NADPH in the presence of Glutamate Dehydrogenase (GDH) to form L-Glutamate and oxidized NADP^+^ and water. The decrease in absorbance at 340 nm, due to the oxidation of NADPH, is proportional to the ammonia concentration.

### Cytosolic pH measurement using pHluorin

Calibration: The strains were grown in media containing 0.1% proline as the sole nitrogen source until the OD_600_ reached 1.0 and were collected and resuspended in PBS buffer containing 100 µg/ml digitonin. This mixture was agitated gently for 15 min using a circular rotor. The cells were pelleted down and resuspended in Na_2_HPO_4_/citrate buffer with varying pH from 4.8 to 8.0. The fluorescence was measured at two excitation wavelengths – 395 nm and 475 nm, with the emission wavelength as 510 nm using Jasco FP 8500 Spectrofluoremeter. A non-linear curve (polynomial with order 2) was plotted for each pH against the fluorescence value I_395/475_.

Sample pH measurement: Cells were grown in media containing 0.1% proline as the sole nitrogen source to reach an optical density of 1.0. The fluorescence was measured at two excitation wavelengths – 395 nm and 475 nm, with the emission wavelength as 510 nm for 10 min (600 sec) after adding 2 mM ammonium or water. The fluorescence value I_395/475_ was calculated by extrapolating from the calibration curve. The pH profile obtained for each strain when water was used as a negative control was subtracted from values obtained when exposed to ammonium. The same analysis was done for triple deletion strain in the presence of water as well as ammonium. ΔpH represents the intracellular pH profile of corresponding strains after subtracting the triple deletion strain values [87].

### FRAP (Fluorescence Recovery After Photobleaching)

Cells were grown in media containing 0.1% proline as the sole nitrogen source for 22 hours until the OD_600_ reaches 3.0. 1 ml of cells were pelleted and collected. Cells were immobilized using 1% agarose pads and were observed using Laser Scanning Microscope (LSM 780 from Carl Zeiss Germany) with 100x magnification. The membrane with yEGFP tagged protein was subjected to FRAP for 60 seconds using an argon ion laser (488 nm). Region of Interest (ROI) is the area where beaching is done at 100% laser intensity for a fixed area at the membrane. The negative control is the membrane area where bleaching is not done. The background is the background fluorescence of the sample slide. Normalized fluorescence intensity was analysed using the double normalization method used in the easyFRAP-web online tool [107][108].

### Plasma membrane isolation

Strains pre-grown in YPD were induced in 0.1% proline medium for 22 hours until the OD_600_ reaches 3.0. Cells were harvested by spinning at 13000 rpm for 15 minutes at 4°C. Cells were washed and resuspended in lysis buffer (50 mM Tris base pH 7.4, 0.5 M HCl, 20% (V/V) Glycerol, 2 mM PMSF, 0.05 mg/ml Protease Inhibitor cocktail) and disrupted using beat beating using glass beads (5 × 1 min with 1 min intervals on ice). The cell lysate was spun at 200,000 g for 90 minutes using a 100 Ti rotor in Beckman Coulter XPN 100 Ultracentrifuge to obtain the plasma membrane fraction as the pellet. The pellet was resuspended in membrane resuspension buffer (50 mM Tris base pH 7.4, 0.2 M HCl, 10% (V/V) Glycerol, 2 mM PMSF, 0.05 mg/ml Protease Inhibitor cocktail)

### Trypsin digestion

The concentration of protein was estimated using SDS PAGE and Pma1 was used as a loading control. The specified concentration of trypsin was added to the membrane fraction containing the desired protein (membrane resuspension buffer did not have PMSF and protein inhibitor cocktail) and was kept at 37°C for 5 minutes. 1 µl of 10 mg/ml trypsin inhibitor after incubation and SDS PAGE was carried out.

## Acknowledgments

Revathi S Iyer was supported by Department of Science and Technology, Government of India WOS-‘A’ Scheme (SR/WOS-A/LS-58/2017). We thank IRCC at Indian Institute of Technology Bombay, India for providing access to the Laser scanning confocal facility. We thank Supreet Saini for support and discussions. We also thank Prof. G. Krishnamoorthy for helping us in designing the FRAP experiments.

## Declaration of interests

The authors declare that they have no known competing financial interests or personal relationships that could have appeared to influence the work reported in this paper.

## Notes

### Competing Interest Statement

The authors have declared no competing interest.

### Summary of Updates

Acknowledgments section is revised

## References

1. K. Bryson, N.J. Msindai, On the Origin of Species by Means of Natural Selection, or the Preservation of Favoured Races in the Struggle for Life, Br. Foreign Medico-Chirurgical Rev. 25 (1860) 367. https://doi.org/10.4324/9781912281244.

2. F. Griffith, The Significance of Pneumococcal Types, J. Hyg. (Lond). 27 (1928) 113–159. https://doi.org/10.1017/S0022172400031879.

3. E.L. Tatum, J. Lederberg, Gene Recombination in the Bacterium Escherichia coli, J. Bacteriol. 53 (1947) 673–684. https://doi.org/10.1128/JB.53.6.673-684.1947.

4. N.D. Zinder, J. Lederberg, Genetic exchange in Salmonella, J. Bacteriol. 64 (1952) 679–699. https://doi.org/10.1128/JB.64.5.679-699.1952.

5. H. Ochman, J.G. Lawrence, E.A. Grolsman, Lateral gene transfer and the nature of bacterial innovation, Nature. 405 (2000) 299–304. https://doi.org/10.1038/35012500.

6. J.D. Elsas, M.J. Bailey, The ecology of transfer of mobile genetic elements, FEMS Microbiol. Ecol. 42 (2002) 187–197. https://doi.org/10.1111/J.1574-6941.2002.TB01008.X.

7. E. V. Koonin, Horizontal gene transfer: essentiality and evolvability in prokaryotes, and roles in evolutionary transitions, F1000Research. 5 (2016). https://doi.org/10.12688/F1000RESEARCH.8737.1.

8. C.G. Kurland, B. Canback, O.G. Berg, Horizontal gene transfer: a critical view, Proc. Natl. Acad. Sci. U. S. A. 100 (2003) 9658–9662. https://doi.org/10.1073/PNAS.1632870100.

9. F. Husnik, J.P. McCutcheon, Functional horizontal gene transfer from bacteria to eukaryotes, Nat. Rev. Microbiol. 16 (2018) 67–79. https://doi.org/10.1038/NRMICRO.2017.137.

10. R. Acuña, B.E. Padilla, C.P. Flórez-Ramos, J.D. Rubio, J.C. Herrera, P. Benavides, S.J. Lee, T.H. Yeats, A.N. Egan, J.J. Doyle, J.K.C. Rose, Adaptive horizontal transfer of a bacterial gene to an invasive insect pest of coffee, Proc. Natl. Acad. Sci. U. S. A. 109 (2012) 4197–4202. https://doi.org/10.1073/PNAS.1121190109.

11. L. Blondel, T.E.M. Jones, C.G. Extavour, Bacterial contribution to genesis of the novel germ line determinant oskar, Elife. 9 (2020). https://doi.org/10.7554/ELIFE.45539.

12. K.B. Sieber, R.E. Bromley, J.C. Dunning Hotopp, Lateral gene transfer between prokaryotes and eukaryotes, Exp. Cell Res. 358 (2017) 421–426. https://doi.org/10.1016/J.YEXCR.2017.02.009.

13. J.C. Dunning Hotopp, M.E. Clark, D.C.S.G. Oliveira, J.M. Foster, P. Fischer, M.C. Muñoz Torres, J.D. Giebel, N. Kumar, N. Ishmael, S. Wang, J. Ingram, R. V. Nene, J. Shepard, J. Tomkins, S. Richards, D.J. Spiro, E. Ghedin, B.E. Slatko, H. Tettelin, J.H. Werren, Widespread lateral gene transfer from intracellular bacteria to multicellular eukaryotes, Science. 317 (2007) 1753–1756. https://doi.org/10.1126/SCIENCE.1142490.

14. J.C. Dunning Hotopp, Horizontal gene transfer between bacteria and animals, Trends Genet. 27 (2011) 157–163. https://doi.org/10.1016/J.TIG.2011.01.005.

15. F. Savory, G. Leonard, T.A. Richards, The role of horizontal gene transfer in the evolution of the oomycetes, PLoS Pathog. 11 (2015). https://doi.org/10.1371/JOURNAL.PPAT.1004805.

16. S.M. Soucy, J. Huang, J.P. Gogarten, Horizontal gene transfer: building the web of life, Nat. Rev. Genet. 16 (2015) 472–482. https://doi.org/10.1038/NRG3962.

17. M. Syvanen, Evolutionary implications of horizontal gene transfer, Annu. Rev. Genet. 46 (2012) 341–358. https://doi.org/10.1146/ANNUREV-GENET-110711-155529.

18. T.A. Richards, A. Monier, A tale of two tardigrades, Proc. Natl. Acad. Sci. U. S. A. 113 (2016) 4892. https://doi.org/10.1073/PNAS.1603862113.

19. E.G.J. Danchin, Lateral gene transfer in eukaryotes: tip of the iceberg or of the ice cube?, BMC Biol. 14 (2016). https://doi.org/10.1186/S12915-016-0330-X.

20. N.A. Moran, T. Jarvik, Lateral transfer of genes from fungi underlies carotenoid production in aphids, Science. 328 (2010) 624–627. https://doi.org/10.1126/SCIENCE.1187113.

21. S. Ohno, Evolution by Gene Duplication, Evol. by Gene Duplic. (1970). https://doi.org/10.1007/978-3-642-86659-3.

22. H.M. Hunsperger, T. Randhawa, R.A. Cattolico, Extensive horizontal gene transfer, duplication, and loss of chlorophyll synthesis genes in the algae, BMC Evol. Biol. 15 (2015). https://doi.org/10.1186/S12862-015-0286-4.

23. E.G.J. Danchin, M.N. Rosso, P. Vieira, J. De Almeida-Engler, P.M. Coutinho, B. Henrissat, P. Abad, Multiple lateral gene transfers and duplications have promoted plant parasitism ability in nematodes, Proc. Natl. Acad. Sci. U. S. A. 107 (2010) 17651–17656. https://doi.org/10.1073/PNAS.1008486107/-/DCSUPPLEMENTAL/SAPP.PDF.

24. T.A. Nguyen, J. Greig, A. Khan, C. Goh, G. Jedd, Evolutionary novelty in gravity sensing through horizontal gene transfer and high-order protein assembly, PLoS Biol. 16 (2018). https://doi.org/10.1371/JOURNAL.PBIO.2004920.

25. A.J. Rivas, A.M. Labella, J.J. Borrego, M.L. Lemos, C.R. Osorio, Evidence for horizontal gene transfer, gene duplication and genetic variation as driving forces of the diversity of haemolytic phenotypes in Photobacterium damselae subsp. damselae, FEMS Microbiol. Lett. 355 (2014) 152–162. https://doi.org/10.1111/1574-6968.12464.

26. V. Daubin, G.J. Szöllősi, Horizontal gene transfer and the history of life, Cold Spring Harb. Perspect. Biol. 8 (2016). https://doi.org/10.1101/cshperspect.a018036.

27. L. Boto, Horizontal gene transfer in evolution: facts and challenges, Proceedings. Biol. Sci. 277 (2010) 819–827. https://doi.org/10.1098/RSPB.2009.1679.

28. M. Syvanen, Cross-species gene transfer; implications for a new theory of evolution, J. Theor. Biol. 112 (1985) 333–343. https://doi.org/10.1016/S0022-5193(85)80291-5.

29. R. Jain, M.C. Rivera, J.E. Moore, J.A. Lake, Horizontal gene transfer accelerates genome innovation and evolution, Mol. Biol. Evol. 20 (2003) 1598–1602. https://doi.org/10.1093/MOLBEV/MSG154.

30. P.R. Marri, G.B. Golding, Gene amelioration demonstrated: the journey of nascent genes in bacteria, Genome. 51 (2008) 164–168. https://doi.org/10.1139/G07-105.

31. A. Medrano-Soto, G. Moreno-Hagelsieb, P. Vinuesa, J.A. Christen, J. Collado-Vides, Successful lateral transfer requires codon usage compatibility between foreign genes and recipient genomes, Mol. Biol. Evol. 21 (2004) 1884–1894. https://doi.org/10.1093/MOLBEV/MSH202.

32. W.W. Navarre, S. Porwollik, Y. Wang, M. McClelland, H. Rosen, S.J. Libby, F.C. Fang, Selective silencing of foreign DNA with low GC content by the H-NS protein in Salmonella, Science. 313 (2006) 236–238. https://doi.org/10.1126/SCIENCE.1128794.

33. T. Kloesges, O. Popa, W. Martin, T. Dagan, Networks of gene sharing among 329 proteobacterial genomes reveal differences in lateral gene transfer frequency at different phylogenetic depths, Mol. Biol. Evol. 28 (2011) 1057–1074. https://doi.org/10.1093/MOLBEV/MSQ297.

34. T. Tuller, Y. Girshovich, Y. Sella, A. Kreimer, S. Freilich, M. Kupiec, U. Gophna, E. Ruppin, Association between translation efficiency and horizontal gene transfer within microbial communities, Nucleic Acids Res. 39 (2011) 4743–4755. https://doi.org/10.1093/NAR/GKR054.

35. G. Kudla, A.W. Murray, D. Tollervey, J.B. Plotkin, Coding-sequence determinants of gene expression in Escherichia coli, Science. 324 (2009) 255–258. https://doi.org/10.1126/SCIENCE.1170160.

36. R. Sorek, Y. Zhu, C.J. Creevey, M.P. Francino, P. Bork, E.M. Rubin, Genome-wide experimental determination of barriers to horizontal gene transfer, Science. 318 (2007) 1449–1452. https://doi.org/10.1126/SCIENCE.1147112.

37. H. Acar Kirit, M. Lagator, J.P. Bollback, Experimental determination of evolutionary barriers to horizontal gene transfer, BMC Microbiol. 20 (2020). https://doi.org/10.1186/S12866-020-01983-5.

38. A. Porse, T.S. Schou, C. Munck, M.M.H. Ellabaan, M.O.A. Sommer, Biochemical mechanisms determine the functional compatibility of heterologous genes, Nat. Commun. 9 (2018). https://doi.org/10.1038/S41467-018-02944-3.

39. O. Popa, T. Dagan, Trends and barriers to lateral gene transfer in prokaryotes, Curr. Opin. Microbiol. 14 (2011) 615–623. https://doi.org/10.1016/J.MIB.2011.07.027.

40. R. Jain, M.C. Rivera, J.A. Lake, Horizontal gene transfer among genomes: The complexity hypothesis, Proc. Natl. Acad. Sci. U. S. A. 96 (1999) 3801–3806. https://doi.org/10.1073/pnas.96.7.3801.

41. O. Cohen, U. Gophna, T. Pupko, The complexity hypothesis revisited: connectivity rather than function constitutes a barrier to horizontal gene transfer, Mol. Biol. Evol. 28 (2011) 1481–1489. https://doi.org/10.1093/MOLBEV/MSQ333.

42. T.A. Richards, G. Leonard, D.M. Soanes, N.J. Talbot, Gene transfer into the fungi, Fungal Biol. Rev. 25 (2011) 98–110. https://doi.org/10.1016/J.FBR.2011.04.003.

43. M. Nagy, F. Lacroute, D. Thomas, Divergent evolution of pyrimidine biosynthesis between anaerobic and aerobic yeasts, Proc. Natl. Acad. Sci. U. S. A. 89 (1992) 8966–8970. https://doi.org/10.1073/PNAS.89.19.8966.

44. Z. Gojković, W. Knecht, E. Zameitat, J. Warneboldt, J.B. Coutelis, Y. Pynyaha, C. Neuveglise, K. Møller, M. Löffler, J. Piškur, Horizontal gene transfer promoted evolution of the ability to propagate under anaerobic conditions in yeasts, Mol. Genet. Genomics. 271 (2004) 387–393. https://doi.org/10.1007/S00438-004-0995-7.

45. C. Hall, S. Brachat, F.S. Dietrich, Contribution of horizontal gene transfer to the evolution of Saccharomyces cerevisiae, Eukaryot. Cell. 4 (2005) 1102–1115. https://doi.org/10.1128/EC.4.6.1102-1115.2005.

46. C. Hall, F.S. Dietrich, The reacquisition of biotin prototrophy in Saccharomyces cerevisiae involved horizontal gene transfer, gene duplication and gene clustering, Genetics. 177 (2007) 2293–2307. https://doi.org/10.1534/GENETICS.107.074963.

47. C. Gonçalves, P. Gonçalves, Multilayered horizontal operon transfers from bacteria reconstruct a thiamine salvage pathway in yeasts, Proc. Natl. Acad. Sci. U. S. A. 116 (2019) 22219–22228. https://doi.org/10.1073/PNAS.1909844116/-/DCSUPPLEMENTAL.

48. D.S. Milner, V. Attah, E. Cook, F. Maguire, F.R. Savory, M. Morrison, C.A. Müller, P.G. Foster, N.J. Talbot, G. Leonard, T.A. Richards, Environment-dependent fitness gains can be driven by horizontal gene transfer of transporter-encoding genes, Proc. Natl. Acad. Sci. U. S. A. 116 (2019) 5613–5622. https://doi.org/10.1073/PNAS.1815994116/-/DCSUPPLEMENTAL.

49. M.A. Coelho, C. Gonçalves, J.P. Sampaio, P. Gonçalves, Extensive intra-kingdom horizontal gene transfer converging on a fungal fructose transporter gene, PLoS Genet. 9 (2013). https://doi.org/10.1371/JOURNAL.PGEN.1003587.

50. M. Novo, F. Bigey, E. Beyne, V. Galeote, F. Gavory, S. Mallet, B. Cambon, J.L. Legras, P. Wincker, S. Casaregola, S. Dequin, Eukaryote-to-eukaryote gene transfer events revealed by the genome sequence of the wine yeast Saccharomyces cerevisiae EC1118, Proc. Natl. Acad. Sci. U. S. A. 106 (2009) 16333–16338. https://doi.org/10.1073/PNAS.0904673106.

51. T.R. McDonald, J.M. Ward, Evolution of electrogenic ammonium transporters (AMTs), Front. Plant Sci. (2016). https://doi.org/10.3389/fpls.2016.00352.

52. J. Peng, C.H. Huang, Rh proteins vs Amt proteins: an organismal and phylogenetic perspective on CO2 and NH3 gas channels, Transfus. Clin. Biol. (2006). https://doi.org/10.1016/j.tracli.2006.02.006.

53. T.R. McDonald, F.S. Dietrich, F. Lutzoni, Multiple horizontal gene transfers of ammonium transporters/ammonia permeases from prokaryotes to eukaryotes: Toward a new functional and evolutionary classification, Mol. Biol. Evol. (2012). https://doi.org/10.1093/molbev/msr123.

54. C.H. Huang, J. Peng, Evolutionary conservation and diversification of Rh family genes and proteins, Proc. Natl. Acad. Sci. U. S. A. (2005). https://doi.org/10.1073/pnas.0507886102.

55. G. Matassi, Horizontal gene transfer drives the evolution of Rh50 permeases in prokaryotes, BMC Evol. Biol. (2017). https://doi.org/10.1186/s12862-016-0850-6.

56. S. Kumar, G. Stecher, M. Suleski, S.B. Hedges, TimeTree: A Resource for Timelines, Timetrees, and Divergence Times, Mol. Biol. Evol. 34 (2017) 1812–1819. https://doi.org/10.1093/MOLBEV/MSX116.

57. I. Eichinger, J.A. Pachebat, G. Glöckner, M.A. Rajandream, R. Sucgang, M. Berriman, J. Song, R. Olsen, K. Szafranski, Q. Xu, B. Tunggal, S. Kummerfeld, M. Madera, B.A. Konfortov, F. Rivero, A.T. Bankier, R. Lehmann, N. Hamlin, R. Davies, P. Gaudet, P. Fey, K. Pilcher, G. Chen, D. Saunders, E. Sodergren, P. Davis, A. Kerhornou, X. Nie, N. Hall, C. Anjard, L. Hemphill, N. Bason, P. Farbrother, B. Desany, E. Just, T. Morio, R. Rost, C. Churcher, J. Cooper, S. Haydock, N. Van Driessche, A. Cronin, I. Goodhead, D. Muzny, T. Mourier, A. Pain, M. Lu, D. Harper, R. Lindsay, H. Hauser, K. James, M. Quiles, M. Madan Babu, T. Saito, C. Buchrieser, A. Wardroper, M. Felder, M. Thangavelu, D. Johnson, A. Knights, H. Loulseged, K. Mungall, K. Oliver, C. Price, M.A. Quail, H. Urushihara, J. Hernandez, E. Rabbinowitsch, D. Steffen, M. Sanders, J. Ma, Y. Kohara, S. Sharp, M. Simmonds, S. Spiegler, A. Tivey, S. Sugano, B. White, D. Walker, J. Woodward, T. Winckler, Y. Tanaka, G. Shaulsky, M. Schleicher, G. Weinstock, A. Rosenthal, E.C. Cox, R.L. Chisholm, R. Gibbs, W.F. Loomis, M. Platzer, R.R. Kay, J. Williams, P.H. Dear, A.A. Noegel, B. Barrell, A. Kuspa, The genome of the social amoeba Dictyostelium discoideum, Nature. 435 (2005) 43–57. https://doi.org/10.1038/nature03481.

58. R. Yoshino, T. Morio, Y. Yamada, H. Kuwayama, M. Sameshima, Y. Tanaka, H. Sesaki, M. Iijima, Regulation of ammonia homeostasis by the ammonium transporter AmtA in Dictyostelium discoideum, Eukaryot. Cell. (2007). https://doi.org/10.1128/EC.00204-07.

59. A.M. Marini, G. Matassi, V. Raynal, B. André, J.P. Cartron, B. Chérif-Zahar, The human Rhesus-associated RhAG protein and a kidney homologue promote ammonium transport in yeast, Nat. Genet. (2000). https://doi.org/10.1038/81656.

60. O. Ninnemann, J.C. Jauniaux, W.B. Frommer, Identification of a high affinity NH4+ transporter from plants, EMBO J. (1994). https://doi.org/10.1002/j.1460-2075.1994.tb06652.x.

61. J. Wang, T. Fulford, Q. Shao, A. Javelle, H. Yang, W. Zhu, M. Merrick, Ammonium transport proteins with changes in one of the conserved pore histidines have different performance in ammonia and methylamine conduction, PLoS One. 8 (2013). https://doi.org/10.1371/JOURNAL.PONE.0062745.

62. A. Adlimoghaddam, M. Boeckstaens, A.M. Marini, J.R. Treberg, A.K.C. Brassinga, D. Weihrauch, Ammonia excretion in Caenorhabditis elegans: Mechanism and evidence of ammonia transport of the Rhesus protein CeRhr-1, J. Exp. Biol. 218 (2015) 675–683. https://doi.org/10.1242/JEB.111856/-/DC1.

63. T.F. Clarke IV, P.L. Clark, Rare codons cluster, PLoS One. 3 (2008) e3412. https://doi.org/10.1371/journal.pone.0003412.

64. A. Rodriguez, G. Wright, S. Emrich, P.L. Clark, %MinMax: A versatile tool for calculating and comparing synonymous codon usage and its impact on protein folding, Protein Sci. 27 (2018) 356. https://doi.org/10.1002/PRO.3336.

65. A.M. Marini, S. Vissers, A. Urrestarazu, B. André, Cloning and expression of the MEP1 gene encoding an ammonium transporter in Saccharomyces cerevisiae, EMBO J. (1994). https://doi.org/10.1002/j.1460-2075.1994.tb06651.x.

66. M. Liutkute, E. Samatova, M. V. Rodnina, Cotranslational Folding of Proteins on the Ribosome, Biomolecules. 10 (2020). https://doi.org/10.3390/BIOM10010097.

67. M. Bogdanov, E. Mileykovskaya, W. Dowhan, Lipids in the Assembly of Membrane Proteins and Organization of Protein Supercomplexes: Implications for Lipid-Linked Disorders, Subcell. Biochem. 49 (2008) 197. https://doi.org/10.1007/978-1-4020-8831-5_8.

68. E. Angov, C.J. Hillier, R.L. Kincaid, J.A. Lyon, Heterologous Protein Expression Is Enhanced by Harmonizing the Codon Usage Frequencies of the Target Gene with those of the Expression Host, PLoS One. 3 (2008) e2189. https://doi.org/10.1371/JOURNAL.PONE.0002189.

69. A.K. Hess, P. Saffert, K. Liebeton, Z. Ignatova, Optimization of Translation Profiles Enhances Protein Expression and Solubility, PLoS One. 10 (2015) e0127039. https://doi.org/10.1371/JOURNAL.PONE.0127039.

70. Z.R. Newman, J.M. Young, N.T. Ingolia, G.M. Barton, Differences in codon bias and GC content contribute to the balanced expression of TLR7 and TLR9, Proc. Natl. Acad. Sci. U. S. A. 113 (2016) E1362–E1371. https://doi.org/10.1073/PNAS.1518976113/-/DCSUPPLEMENTAL.

71. N. Behloul, W. Wei, S. Baha, Z. Liu, J. Wen, J. Meng, Effects of mRNA secondary structure on the expression of HEV ORF2 proteins in Escherichia coli, Microb. Cell Fact. 16 (2017). https://doi.org/10.1186/S12934-017-0812-8.

72. A.M. Marini, M. Boeckstaens, F. Benjelloun, B. Chérif-Zahar, B. André, Structural involvement in substrate recognition of an essential aspartate residue conserved in Mep/Amt and Rh-type ammonium transporters, Curr. Genet. 49 (2006) 364–374. https://doi.org/10.1007/S00294-006-0062-5/FIGURES/9.

73. A. Karginov, M. Agaphonov, A simple enrichment procedure improves detection of membrane proteins by immunoblotting, Biotechniques. 61 (2016) 260–261. https://doi.org/10.2144/000114474.

74. A. Rath, M. Glibowicka, V.G. Nadeau, G. Chen, C.M. Deber, Detergent binding explains anomalous SDS-PAGE migration of membrane proteins, Proc. Natl. Acad. Sci. U. S. A. (2009). https://doi.org/10.1073/pnas.0813167106.

75. F. Buhr, S. Jha, M. Thommen, J. Mittelstaet, F. Kutz, H. Schwalbe, M. V. Rodnina, A.A. Komar, Synonymous Codons Direct Cotranslational Folding toward Different Protein Conformations, Mol. Cell. 61 (2016) 341–351. https://doi.org/10.1016/J.MOLCEL.2016.01.008.

76. D. Blakey, A. Leech, G.H. Thomas, G. Coutts, K. Findlay, M. Merrick, Purification of the Escherichia coli ammonium transporter AmtB reveals a trimeric stoichiometry, Biochem. J. (2002). https://doi.org/10.1042/BJ20011761.

77. M.J. Conroy, S.J. Jamieson, D. Blakey, T. Kaufmann, A. Engel, D. Fotiadis, M. Merrick, P.A. Bullough, Electron and atomic force microscopy of the trimeric ammonium transporter AmtB, EMBO Rep. (2004). https://doi.org/10.1038/sj.embor.7400296.

78. S.H. Bokman, W.W. Ward, Renaturation of Aequorea green-fluorescent protein, Biochem. Biophys. Res. Commun. 101 (1981) 1372–1380. https://doi.org/10.1016/0006-291X(81)91599-0.

79. J.A. Sniegowski, M.E. Phail, R.M. Wachter, Maturation efficiency, trypsin sensitivity, and optical properties of Arg96, Glu222, and Gly67 variants of green fluorescent protein, Biochem. Biophys. Res. Commun. 332 (2005) 657–663. https://doi.org/10.1016/J.BBRC.2005.04.166.

80. J.S. Reuter, D.H. Mathews, RNAstructure: Software for RNA secondary structure prediction and analysis, BMC Bioinformatics. (2010). https://doi.org/10.1186/1471-2105-11-129.

81. C. Kimchi-Sarfaty, J.M. Oh, I.W. Kim, Z.E. Sauna, A.M. Calcagno, S. V. Ambudkar, M.M. Gottesman, A “silent” polymorphism in the MDR1 gene changes substrate specificity, Science (80-.). (2007). https://doi.org/10.1126/science.1135308.

82. R.C. Hunt, V.L. Simhadri, M. Iandoli, Z.E. Sauna, C. Kimchi-Sarfaty, Exposing synonymous mutations, Trends Genet. 30 (2014) 308–321. https://doi.org/10.1016/J.TIG.2014.04.006.

83. S. Kirchner, Z. Cai, R. Rauscher, N. Kastelic, M. Anding, A. Czech, B. Kleizen, L.S. Ostedgaard, I. Braakman, D.N. Sheppard, Z. Ignatova, Alteration of protein function by a silent polymorphism linked to tRNA abundance, PLOS Biol. 15 (2017) e2000779. https://doi.org/10.1371/JOURNAL.PBIO.2000779.

84. M. Boeckstaens, B. André, A.M. Marini, Distinct transport mechanisms in yeast ammonium transport/sensor proteins of the Mep/Amt/Rh family and impact on filamentation, J. Biol. Chem. (2008). https://doi.org/10.1074/jbc.M801467200.

85. M. Boeckstaens, B. André, A.M. Marini, The yeast ammonium transport protein Mep2 and its positive regulator, the Npr1 kinase, play an important role in normal and pseudohyphal growth on various nitrogen media through retrieval of excreted ammonium, Mol. Microbiol. 64 (2007) 534–546. https://doi.org/10.1111/J.1365-2958.2007.05681.X.

86. E. Dubois, M. Grenson, Methylamine/ammonia uptake systems in Saccharomyces cerevisiae: multiplicity and regulation, MGG Mol. Gen. Genet. (1979). https://doi.org/10.1007/BF00267857.

87. A.S. Brito, B. Neuhauser, R. Wintjens, A.M. Marini, M. Boeckstaens, Yeast filamentation signaling is connected to a specific substrate translocation mechanism of the Mep2 transceptor, PLoS Genet. (2020). https://doi.org/10.1371/journal.pgen.1008634.

88. S. Khademi, J. O’Connell, J. Remis, Y. Robles-Colmenares, L.J.W. Miercke, R.M. Stroud, Mechanism of ammonia transport by Amt/MEP/Rh: Structure of AmtB at 135 Å, Science (80-.). 305 (2004) 1587–1594. https://doi.org/10.1126/science.1101952.

89. T. Wacker, J.J. Garcia-Celma, P. Lewe, S.L.A. Andrade, Direct observation of electrogenic NH4+ transport in ammonium transport (Amt) proteins, Proc. Natl. Acad. Sci. U. S. A. 111 (2014) 9995–10000. https://doi.org/10.1073/pnas.1406409111.

90. G. Williamson, G. Tamburrino, A. Bizior, M. Boeckstaens, G.D. Mirandela, M. Bage, A. Pisliakov, C.M. Ives, E. Terras, P.A. Hoskisson, A.M. Marini, U. Zachariae, A. Javelle, A two-lane mechanism for selective biological ammonium transport, Elife. 9 (2020) 1–41. https://doi.org/10.7554/eLife.57183.

91. T.N. Starr, J.W. Thornton, Epistasis in protein evolution, Protein Sci. 25 (2016) 1204–1218. https://doi.org/10.1002/PRO.2897.

92. F.J. Poelwijk, M. Socolich, R. Ranganathan, Learning the pattern of epistasis linking genotype and phenotype in a protein, Nat. Commun. 2019 101. 10 (2019) 1–11. https://doi.org/10.1038/s41467-019-12130-8.

93. C.E. Gonzalez, M. Ostermeier, Pervasive Pairwise Intragenic Epistasis among Sequential Mutations in TEM-1 β-Lactamase, J. Mol. Biol. 431 (2019) 1981–1992. https://doi.org/10.1016/J.JMB.2019.03.020.

94. J.R. Iben, R.J. Maraia, tRNAomics: tRNA gene copy number variation and codon use provide bioinformatic evidence of a new anticodon:codon wobble pair in a eukaryote, RNA. 18 (2012) 1358. https://doi.org/10.1261/RNA.032151.111.

95. E. Zuckerkandl, L. Pauling, Molecules as documents of history, J. Theor. Biol. 8 (1965) 357–366.

96. C. Yanofsky, V. Horn, D. Thorpe, Protein structure relationships revealed by mutational analysis, Science (80-.). (1964). https://doi.org/10.1126/science.146.3651.1593.

97. B.A. Malcolm, K.P. Wilson, B.W. Matthews, J.F. Kirsch, A.C. Wilson, Ancestral lysozymes reconstructed, neutrality tested, and thermostability linked to hydrocarbon packing, Nature. 345 (1990) 86–89. https://doi.org/10.1038/345086a0.

98. L. Tubiana, A.L. Božič, C. Micheletti, R. Podgornik, Synonymous mutations reduce genome compactness in icosahedral ssRNA viruses, Biophys. J. 108 (2015) 194–202. https://doi.org/10.1016/J.BPJ.2014.10.070/ATTACHMENT/11C1A729-E23F-4336-AFF5-FCAF1387C8D3/MMC1.PDF.

99. M.P. Zwart, M.F. Schenk, S. Hwang, B. Koopmanschap, N. de Lange, L. van de Pol, T.T.T. Nga, I.G. Szendro, J. Krug, J.A.G.M. de Visser, Unraveling the causes of adaptive benefits of synonymous mutations in TEM-1 β-lactamase, Heredity (Edinb). (2018). https://doi.org/10.1038/s41437-018-0104-z.

100. J. Duan, M.S. Wainwright, J.M. Comeron, N. Saitou, A.R. Sanders, J. Gelernter, P. V. Gejman, Synonymous mutations in the human dopamine receptor D2 (DRD2) affect mRNA stability and synthesis of the receptor, Hum. Mol. Genet. 12 (2003) 205–216. https://doi.org/10.1093/HMG/DDG055.

101. S.A. Shabalina, N.A. Spiridonov, A. Kashina, Sounds of silence: synonymous nucleotides as a key to biological regulation and complexity, Nucleic Acids Res. 41 (2013) 2073–2094. https://doi.org/10.1093/NAR/GKS1205.

102. V. Bali, A. Lazrak, P. Guroji, L. Fu, S. Matalon, Z. Bebok, A synonymous codon change alters the drug sensitivity of ΔF508 cystic fibrosis transmembrane conductance regulator, FASEB J. 30 (2016) 201–213. https://doi.org/10.1096/FJ.15-273714.

103. Y. Sharma, M. Miladi, S. Dukare, K. Boulay, M. Caudron-Herger, M. Groß, R. Backofen, S. Diederichs, A pan-cancer analysis of synonymous mutations, Nat. Commun. 10 (2019) 1–14. https://doi.org/10.1038/s41467-019-10489-2.

104. V. Gotea, J.J. Gartner, N. Qutob, L. Elnitski, Y. Samuels, The functional relevance of somatic synonymous mutations in melanoma and other cancers, Pigment Cell Melanoma Res, 2015. https://doi.org/10.1111/pcmr.12413.

105. K. Vetsigian, C. Woese, N. Goldenfeld, Collective evolution and the genetic code, Proc. Natl. Acad. Sci. U. S. A. 103 (2006) 10696–10701. https://doi.org/10.1073/PNAS.0603780103.

106. Molecular Cloning: A Laboratory Manual, 4th edition., 2012. www.cshlpress.org.

107. easyFRAP-web | Online FRAP Analysis Tool, (n.d.). https://easyfrap.vmnet.upatras.gr/?AspxAutoDetectCookieSupport=1 (accessed March 2, 2022).

108. G. Koulouras, A. Panagopoulos, M.A. Rapsomaniki, N.N. Giakoumakis, S. Taraviras, Z. Lygerou, EasyFRAP-web: a web-based tool for the analysis of fluorescence recovery after photobleaching data, Nucleic Acids Res. 46 (2018) W467–W472. https://doi.org/10.1093/NAR/GKY508.

