## Supplementary figures for "Epistasis between synonymous and nonsynonymous mutations in *Dictyostelium discoideum* ammonium transporter *amtA* drives functional complementation in *Saccharomyces cerevisiae*"

Figure S1: Growth pattern of *amtA* and *MEP2* on low ammonium at a single colony level

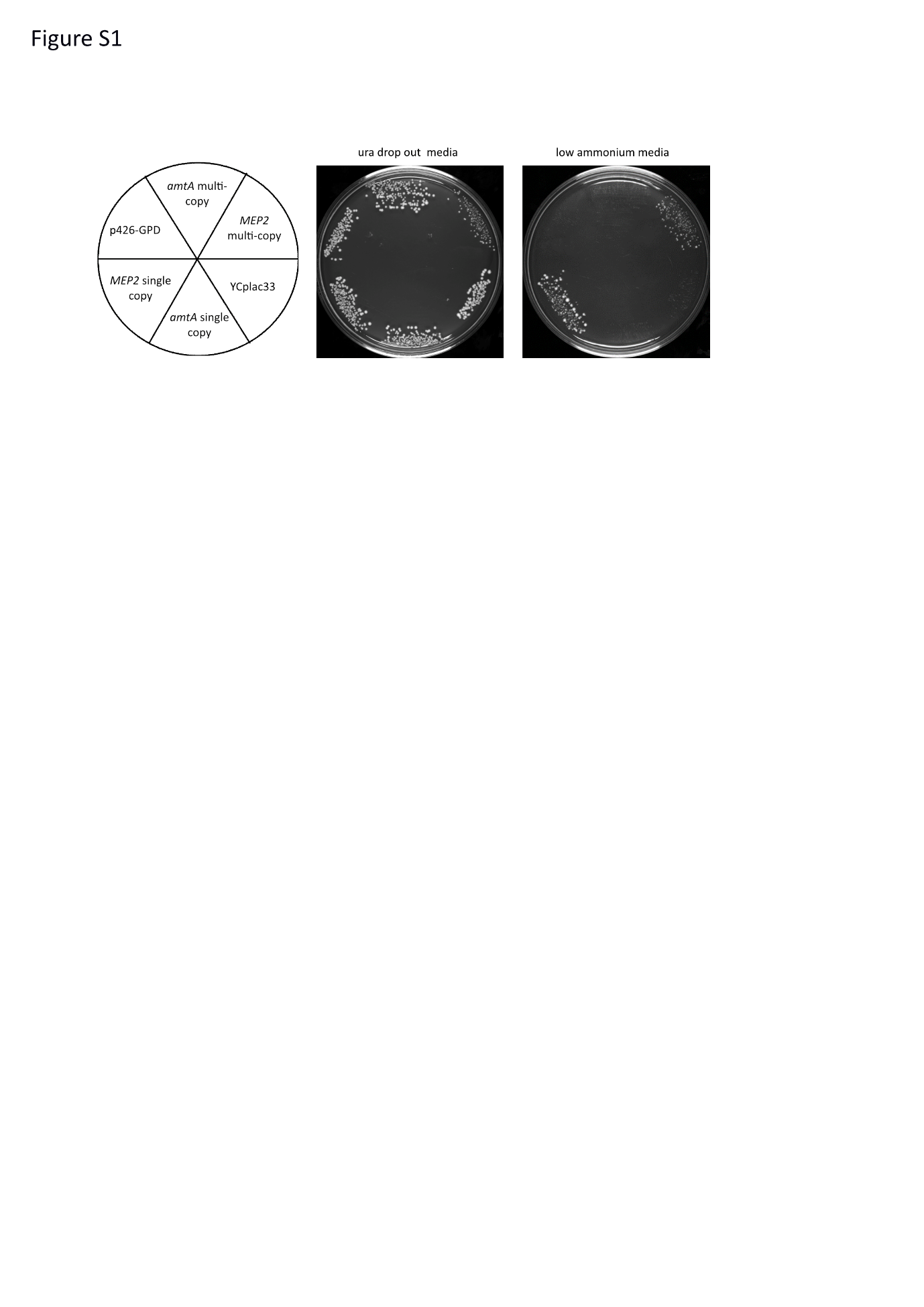

Growth of transformants on low ammonium**.** 5 µl of the transformants of the triple deletion strain pre-grown in ura drop out glucose medium containing 20 mM ammonium to a cell density of 10^7^/ml were streaked onto ura drop out glucose medium with 20 mM ammonium sulfate (middle panel) and low ammonium media (right panel). The growth pattern was photographed after 4 days of incubation at 30ºC. The left panel is a schematic representation of the transformants bearing the corresponding plasmids.

Figure S2: Schematic representation of generation of *amtA* mutant library

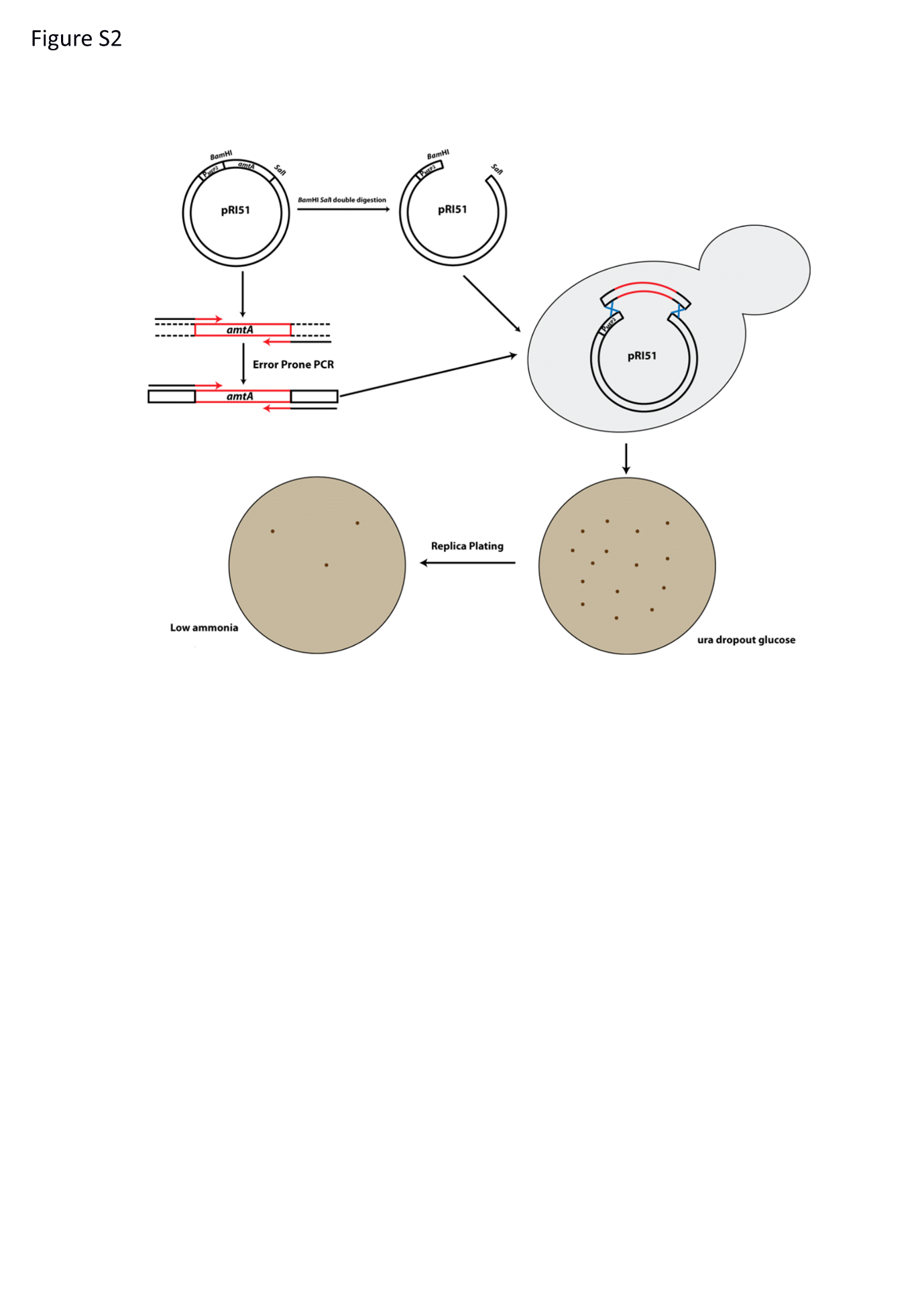

Figure S3: Growth pattern of *amtA* mutants on low ammonium at a single colony level

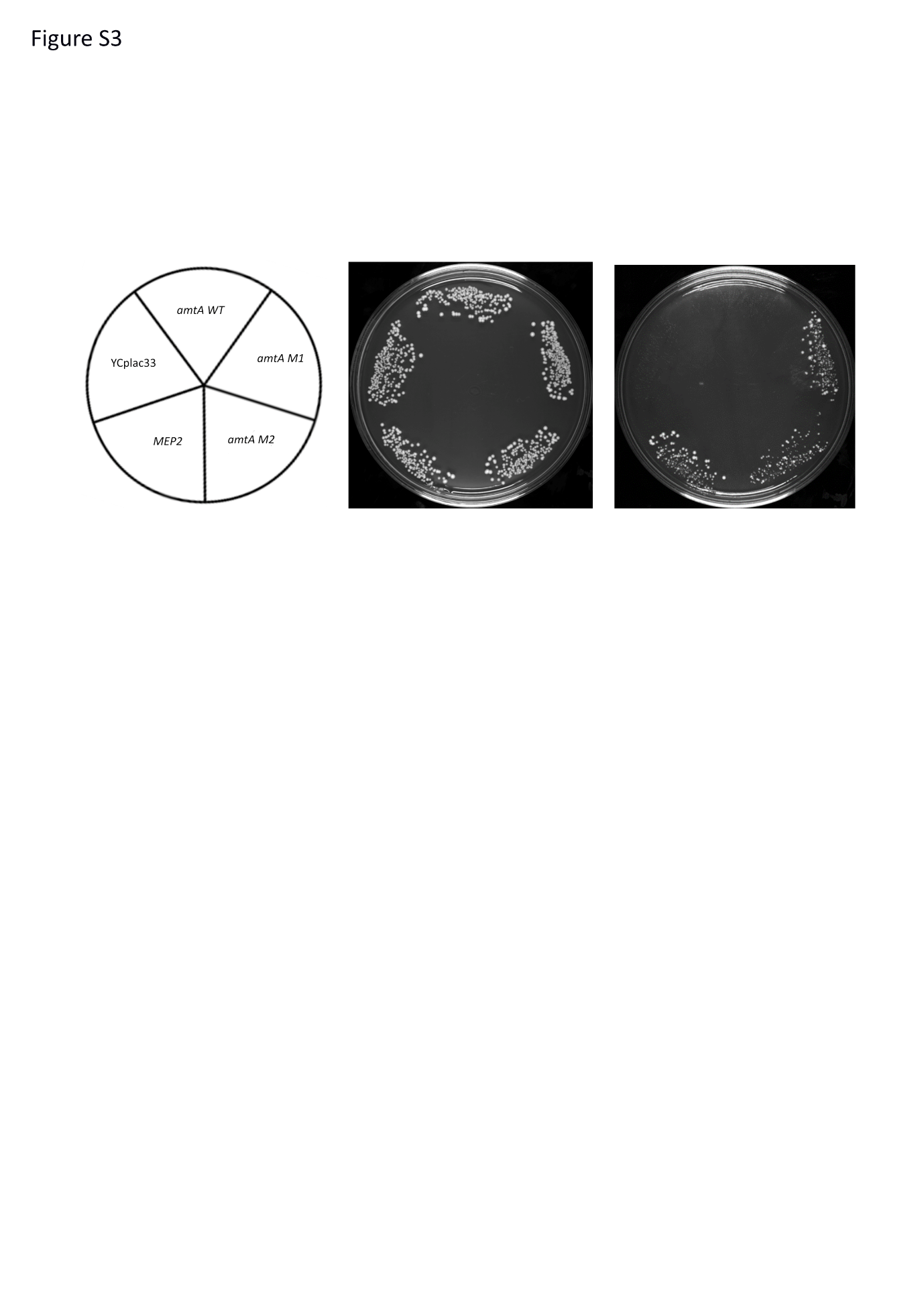

Growth of transformants on low ammonium**.** 5 µl of the transformants of the triple deletion strain pre-grown in ura drop out glucose medium containing 20 mM ammonium to a cell density of 10^7^/ml were streaked onto ura drop out glucose medium with 20 mM ammonium sulfate (middle panel) and low ammonium media (right panel). The growth pattern was photographed after 4 days of incubation at 30ºC. The left panel is a schematic representation of the transformants bearing the corresponding plasmids.

Figure S4: Growth pattern of *yEGFP* tagged strains

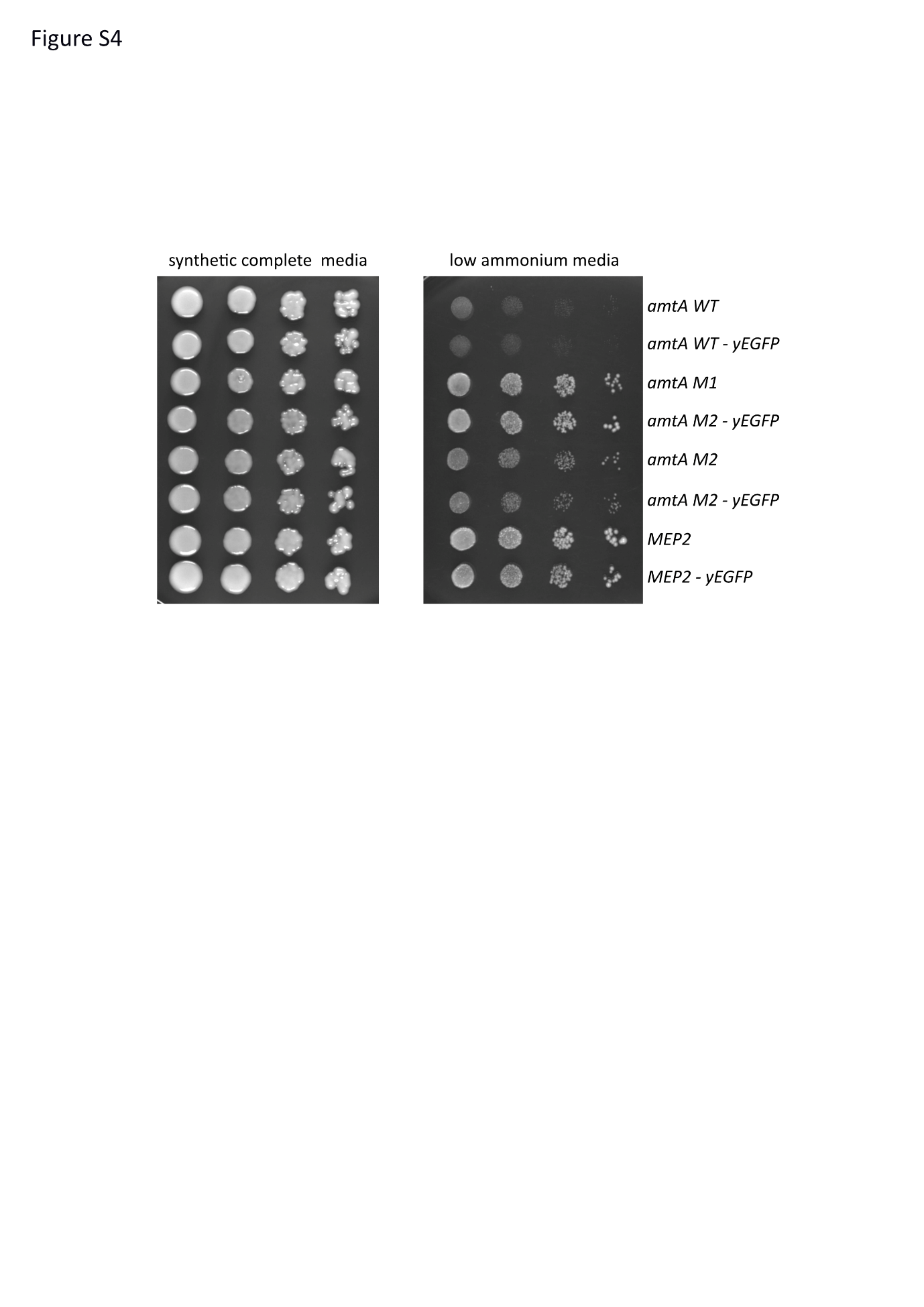

**Growth of strains on low ammonium**. 5 µl of the strains pre-grown in synthetic complete glucose medium containing 20 mM ammonium to a cell density of 10^7^/ml were serially diluted and were spotted onto synthetic complete glucose medium with 20 mM ammonium sulfate (left panel) and low ammonium media (right panel). The growth pattern was photographed after 4 days of incubation at 30ºC.
